# Transcriptional integration of distinct microbial and nutritional signals by the small intestinal epithelium

**DOI:** 10.1101/2021.11.01.465976

**Authors:** Colin R. Lickwar, James M. Davison, Cecelia Kelly, Gilberto Padilla Mercado, Jia Wen, Briana R. Davis, Matthew C. Tillman, Ivana Semova, Sarah F. Andres, Goncalo Vale, Jeffrey G. McDonald, John F. Rawls

## Abstract

To preserve its physiologic functions, the intestine must interpret and adapt to complex combinations of stimuli from dietary and microbial sources. However, the transcriptional strategies by which the intestinal epithelium integrates and adapts to dietary and microbial information remains unresolved. We compared adult mice reared germ free (GF) or conventionalized with a microbiota (CV) either fed normally or after a single high-fat meal (HFM). Jejunal epithelium preparations were queried using genomewide assays for RNA-seq, the activating histone mark H3K27ac ChIP-seq, and ChIP-seq of the microbially-responsive transcription factor HNF4A. We identified distinct nutritional and microbial responses at certain genes, but also apparent simultaneous influence of both stimuli at many other loci and regulatory regions. Increased expression levels and H3K27ac enrichment following HFM at a subset of these sites was dependent on microbial status. H3K27ac sites that were preferentially increased by HFM in the presence of microbes neighbor lipid anabolism and proliferation genes as well as intestinal stem cell (ISC) markers, were usually active only in ISCs, and were not HNF4A targets. In contrast, H3K27ac sites that were preferentially increased by HFM in the absence of microbes neighbored targets of the nuclear receptor and energy homeostasis regulator PPARA, were frequently accessible only in enterocytes, and were HNF4A bound. These results reveal that HNF4A supports a differentiated enterocyte and FAO program in GF, and that suppression of HNF4A by the combination of microbes and HFM may result in preferential activation of IEC proliferation programs. Microbial and nutritional responses are therefore integrated with some of the same transcriptional programs that regulate intestinal proliferation and differentiation.

## Introduction

The intestine simultaneously serves as the primary site for dietary nutrient absorption and as a habitat for resident microbiota. Microbial and nutritional signals are diverse, dynamic, and frequently simultaneous within the gut (Soderholm and Pedicord 2019). However, host diet can dramatically influence microbial communities and their metabolites (Martinez-Guryn et al. 2018; Patnode et al. 2019). Conversely, microbes can modify nutritional signals (Krautkramer et al. 2021). This allows microbes to influence existing host signaling pathways to modulate disease pathogenesis or create beneficial symbioses with the host (Byndloss et al. 2017; Krautkramer et al. 2021; Salvi and Cowles 2021).

Understanding how the intestine perceives and responds to the major stimuli of nutritional and microbial signals remains a fundamental challenge. Frequently, studies interrogate the effect of nutrients or microbes separately without exploring integrative host responses. For example, elevated levels of dietary fat have been shown to exert a dominant effect on energy intake and adiposity in mice (Hu et al. 2018) and are implicated in the global prevalence of human metabolic disorders (Oakes et al. 1997; Ludwig et al. 2018). However, there is ample evidence that high-fat diet and microbiota interactively influence host physiology. For example, germ-free mice are resistant to obesity from a high-fat diet (Backhed et al. 2004; Backhed et al. 2007; Turnbaugh et al. 2008; Martinez-Guryn et al. 2018).

Whereas chronic high-fat diet feeding leads to adaptive physiological responses that can make it difficult to distinguish primary impacts of microbiota on host response (Beyaz et al. 2016; Duan et al. 2018), those impacts can be more easily discerned in the postprandial response to a single high-fat meal (HFM). Complementary studies in zebrafish and mice given a single HFM challenge have established that microbiota colonization promotes dietary fat absorption in intestinal epithelial cells (IECs) and distribution to the rest of the body (Semova et al. 2012; Falcinelli et al. 2015; Martinez-Guryn et al. 2018; Sheng et al. 2018). In both mice and zebrafish, different microbes appear to have contradictory effects on lipid metabolism in IECs, implicating multiple pathways in the integration of these environments (Semova et al. 2012; Falcinelli et al. 2015; Tazi et al. 2018; Araujo et al. 2020). In support, *in vitro* exposure of mouse enteroids or an intestinal epithelial cell line to several bacterial strains and their products have distinct effects on fatty acid metabolism and expression of associated genes (Martinez-Guryn et al. 2018; Tazi et al. 2018; Araujo et al. 2020). However, the mechanisms and pathways underlying these phenotypes *in vivo* remain unknown. Also, how genomewide transcriptional responses in IECs integrate multiple signals simultaneously from a complex microbial community and diet remain unmapped.

Postprandial uptake of dietary lipids takes place primarily in the jejunum region of the small intestine. Genomewide analyses of the intestinal response to microbes consistently show a reduction in expression of lipid metabolism genes in small intestinal IECs and have implicated circadian rhythm, numerous transcription factors (TFs) and other regulators in the response including, *Nfil3, Hdac3*, and the lipid-liganded nuclear receptor TFs including the Ppara and *Hnf4a* (El Aidy et al. 2013; Mukherji et al. 2013; Camp et al. 2014; Davison et al. 2017; Wang et al. 2017; Kuang et al. 2019; Wu et al. 2020). PPARA functions as a major regulator of fatty acid oxidation (FAO) genes and energy homeostasis partly through activating lipid metabolism genes when in the presence of lipids (Grygiel-Gorniak 2014). HNF4A is also a major regulator of lipid metabolism genes in the intestine and liver (Archer et al. 2005; Yin et al. 2011; Frochot et al. 2012; Davison et al. 2017; Qin et al. 2018; Chen et al. 2020). In zebrafish, the majority of microbially suppressed genes, including many lipid metabolism genes, also lose expression in *hnf4a* mutants (Davison et al. 2017). In mouse, colonization results in a reduction of intestinal HNF4A occupancy at most sites across the genome (Davison et al. 2017). Collectively, this suggests HNF4A binding activity and function are high in a germ-free context in IECs and may regulate the response to microbial and nutritional signals. However, the underlying reason for the overlap between microbial and *hnf4a* regulated genes and how microbes alter HNF4A occupancy, host metabolism, and acquisition of nutrients remain unknown.

Here we applied multiple functional genomic assays to evaluate the interaction between high-fat meal (HFM) and microbiota colonization in mouse small intestinal epithelial cells. Entry of a HFM into the small intestine initiates a dynamic postprandial process beginning with emulsification and digestion within the lumen, leading to uptake of lipids into IECs where they are either temporarily stored in cytosolic lipid droplets, oxidized as a fuel source, or exported in chylomicron lipoproteins that are distributed via the circulatory system. We focused here on a single, early postprandial time point following gavage of a complex HFM consisting of chicken egg yolk emulsion, chosen to capture transcriptional responses during the initial perception and response to a complex HFM before cell-division and cell-type changes are substantial. Collectively, our results suggest that microbial and nutritional signal responses are inherently tied to some of the same transcriptional programs that regulate intestinal proliferation and differentiation.

## Results

### Transcriptional changes integrate microbial and nutritional responses in the intestine

By manipulating the presence of microbiota and HFM, we queried four conditions in the adult mouse jejunum: Germ-free (GF), Germ-free plus high-fat meal (GF+HFM), ex-GF colonized with a conventional microbiota for two weeks (Colonized, CV), and Colonized plus high-fat meal (CV+HFM). **(Fig. 1A**). Previous studies suggested that 2h after HFM gavage was sufficient to initiate lipid droplet accumulation and chylomicron export in IECs, but too early in the postprandial process to detect major differences in intestinal lipid transport between GF and CV mice (Backhed et al. 2007; Zhu et al. 2009; Martinez-Guryn et al. 2018). In accord, we found that gavaging mice with HFM consisting of a chicken egg yolk emulsion labeled with BODIPY-conjugated C12 fatty acid led to BODIPY accumulation in jejunal IECs after 2h (**Fig. 1B**). Wholemount confocal microscopy of villi from mice in each +HFM condition identified a trending increase in epithelial BODIPY signal in CV+HFM relative to GF+HFM (**Fig. 1C**). This is consistent with previous reports that microbial colonization promotes lipid accumulation in IECs (Semova et al. 2012; Martinez-Guryn et al. 2018; Tazi et al. 2018) and confirms that lipid has reached IECs by this timepoint. We next performed direct infusion MS/MS^ALL^ lipidomic analysis (Vale et al. 2019) on jejunal IEC preparations (see Supplemental Materials for preparation methods) to assess potential differences in lipid content under these four conditions. We observed no significant differences between the 4 conditions in the relative abundance of major lipid classes but noted a trending increase in triacylglycerol (TAG) in CV and CV+HFM conditions (**Supplemental Fig. S1A; Supplemental Table S1**). Analysis of fatty acid (FA) saturation within neutral lipid pools revealed relative increases in the abundance of saturated FA in both HFM conditions and relative decreases in polyunsaturated FA (**Supplemental Fig. S1B**), presumably reflecting the influx of saturated FA known to predominate in chicken egg yolk (Wood et al. 2021). Closer inspection of individual FA species within the neutral lipid pool confirmed similar increases in 16:0 and 18:0 and reduction in 18:2 in both HFM conditions compared to their controls (**Supplemental Fig. S1C**). Analysis of other FA species and broader lipid species suggested other potential impacts of microbiota and HFM feeding (**Supplemental Fig. S1C-E**). For example, GF mice displayed elevated anti-inflammatory 22:6 (docosahexaenoic acid) relative to CV under both dietary conditions (**Supplemental Fig. S1C**) consistent with a previous study (Liebisch et al. 2021). Together these results establish that 2h of HFM feeding is sufficient to initially incorporate dietary lipids into IECs, with only nominal differences in lipid content between GF and CV states at this early postprandial stage.

**Figure 1:**
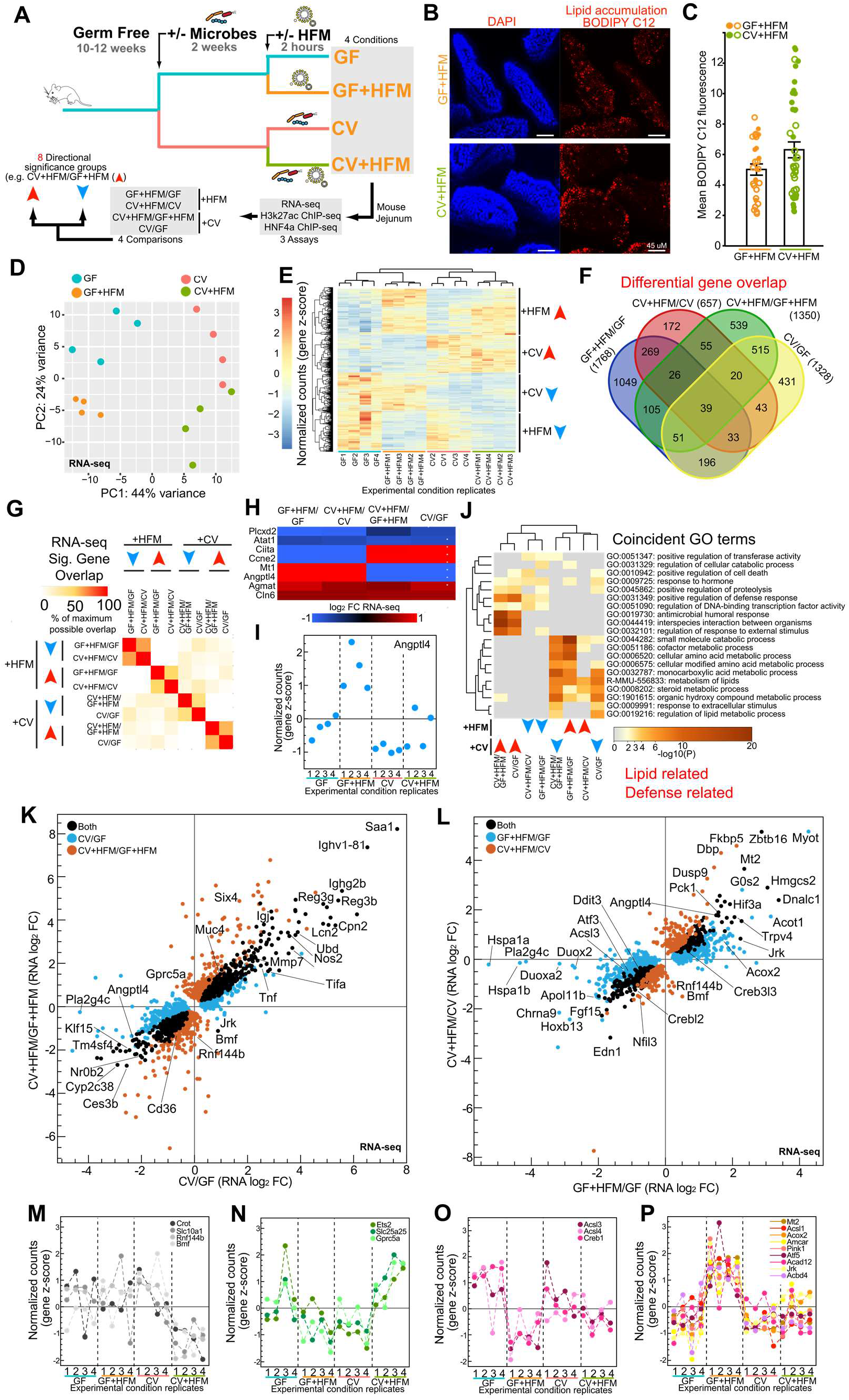
Impact of high fat meal and microbes on the mammalian intestine. **(A)** Experimental schematic highlighting microbial and nutritional conditions, genomic assays, and analysis. **(B)** Confocal *en face* images of jejunal villi 2h post BODIPY C12 egg yolk gavage (red) with nuclei labeled with DAPI (blue). **(C)** Quantification of mean BODIPY C12 fluorescence per villus. Data points represent individual villi (27 GF+HFM villi, 42 CV+HFM villi), with open and closed circles representing villi from two biological replicate mice per condition. Averages and standard deviations of villi measurements are shown. Student’s t-test showed significant differences when comparing across villi (p=0.023) but not mice (p=0.304). **(D)** PCA of RNA-seq normalized counts in each replicate for the 4 conditions. **(E)** Heatmap of row z-scored normalized counts for genes significantly differential in at least one comparison by RNA-seq. Examples of blocks of commonly behaving genes are marked for +HFM and +CV directional groups. **(F)** Venn diagram showing differential genes that overlap across different comparisons. **(G)** Pairwise comparison of maximum overlap of coincident significant RNA-seq genes for 8 directional significance groups show generally coincident directionality and genes for +CV and +HFM comparisons. **(H)** Heatmap of example genes significantly different in both a +CV and +HFM comparison. **(I)** RNA-seq z-scored normalized counts of *Angptl4*, which is significant in both +CV and +HFM comparisons, shows amplified relative expression in the GF+HFM conditions. **(J)** Clustered heatmap of significance values for shared GO terms in at least 2 of 8 directional RNA-seq significance groups. **(K)** Scatterplot of significantly different gene log_2_ fold change for CV/GF and CV+HFM/GF+HFM RNA-seq. **(L)** Same as (**K**) for GF+HFM/GF and CV+HFM/CV. (**M-P**) RNA-seq z-scored normalized counts for example interacting genes.

RNA-seq of jejunal IEC preparations (see Supplemental Materials for preparation methods) showed separation of the four conditions with each experimental replicate clustering based on treatment **(Fig. 1D)**. We identified hundreds of genes with significant transcriptional differences for each of the separate four comparisons (+CV: CV/GF and CV+HFM/GF+HFM and +HFM: GF+HFM/GF and CV+HFM/CV) (**Fig. 1A; Supplemental Table S2**) and identified blocks of genes that were primarily impacted by microbial or nutritional status (**Fig. 1E**). Overlap of differential genes within both +HFM comparisons and both +CV comparisons identified substantial agreement (**Fig. 1F**). However, several genes were also significantly different in response to both +CV and +HFM conditions (**Fig. 1F,G; Supplemental Fig. 2A**; e.g. GF+HFM/GF and CV/GF). As a result, varied patterns of differential expression were evident across comparisons, suggesting +CV and +HFM conditions were integrated at a number of genes (**Fig. 1H; Supplemental Fig. 2B**). These genes include *Angptl4*, a secreted lipoprotein lipase inhibitor involved in partitioning triglyceride availability and lipid accumulation known to be suppressed by microbiota (Backhed et al. 2004; Backhed et al. 2007; Camp et al. 2012; Davison et al. 2017) and activated by fasting and high-fat diet (Kersten et al. 2000; Mattijssen et al. 2014). *Angptl4* showed significant transcriptional differences in all four comparisons (**Fig. 1H,I; Supplemental Fig. 2C**). Interestingly, despite being elevated in GF relative to CV, *Angptl4* is further upregulated by HFM in both GF+HFM and CV+HFM conditions. This increase occurs proportionally such that *Angptl4* is still significantly higher in GF+HFM relative to CV+HFM. This suggests that the nutritional and microbial signals that regulate gene transcription, while separable, can be additive or subtractive in their contributions to regulation of transcription within intestinal epithelia (**Fig. 1I**).

**Figure 2:**
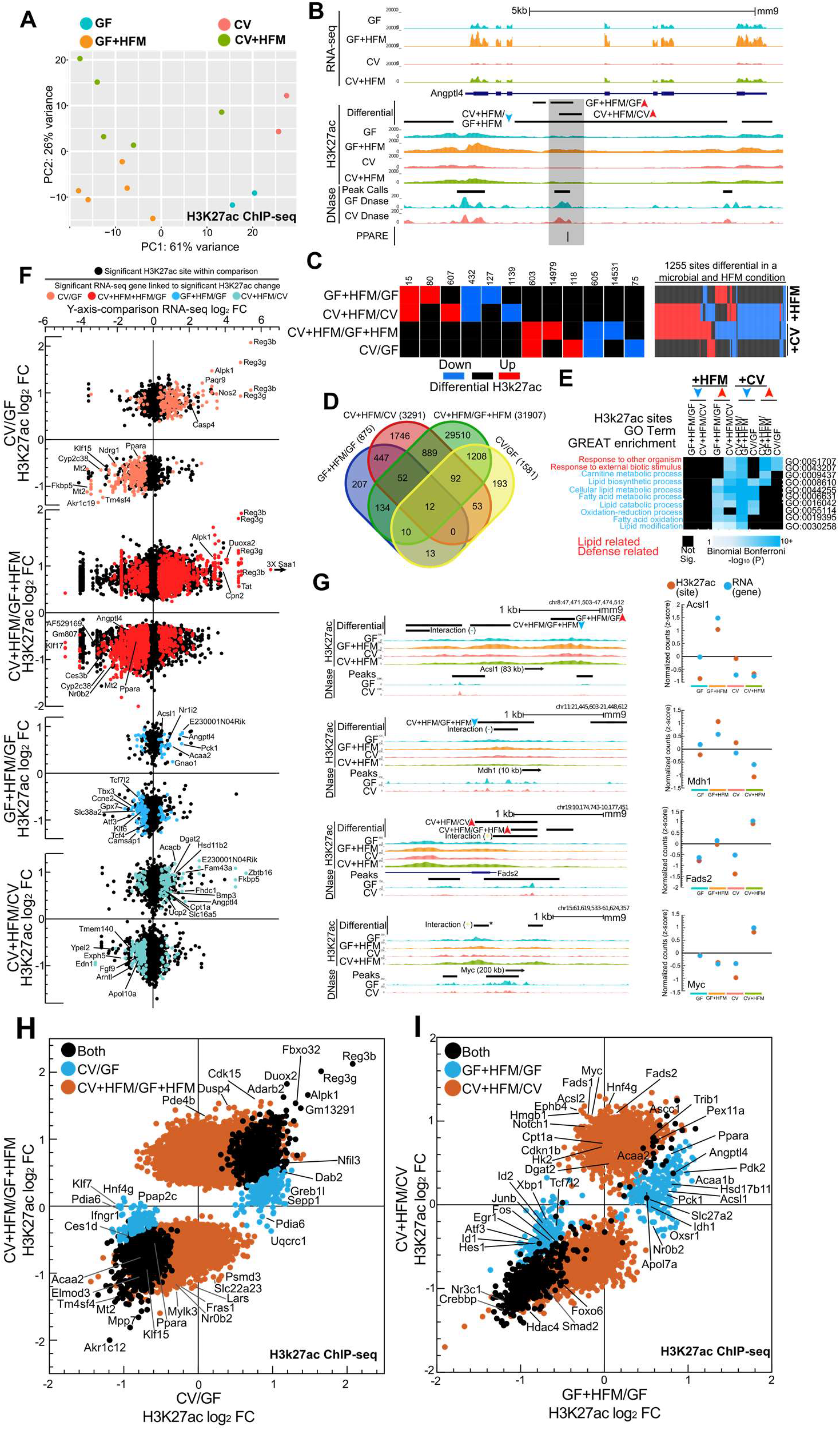
Identification of nutritional and microbial regulatory regions in the mammalian intestine. **(A)** PCA of H3K27ac ChIP-seq normalized counts for all replicates for each condition. **(B)** Average RNA-seq, H3K27ac ChIP-seq, and DNase-seq signal for various conditions at the *Angptl4* locus. An accessible chromatin region coincident with a characterized PPAR binding site in intron 3 was microbially suppressed and +HFM induced (Mandard et al. 2004). **(C)** Quantification of different patterns of differential H3K27ac DNase sites. 1255 regulatory regions are responsive to both +CV and +HFM conditions. **(D)** Venn diagram of overlap for H3K27ac sites for +CV and +HFM comparisons. **(E)** Coincident GREAT GO terms enrichment for 8 H3K27ac directional significance groups (McLean et al. 2010). **(F)** Scatterplots of RNA-seq versus H3K27ac log_2_ fold change for all 4 comparisons. Colored dots represent associated genes significant by RNA-seq in that condition and associated with H3K27ac sites. **(G)** H3K27ac ChIP-seq signal at H3K27ac interaction regulatory regions that share common patterns of H3K27ac signal and RNA across all four (right). Due to two sites, the represented *Myc* interaction site is marked with an asterisk. **(H)** Scatterplot of average H3K27ac sites with a significantly different log_2_ fold change window for CV/GF and CV+HFM/GF+HFM. **(I)** Same as (H) for GF+HFM/GF and CV+HFM/CV.

We next used Gene Ontology (GO) terms to functionally categorize each of the comparisons broken down into genes that were up- or down-regulated to create 8 total differential significance groups (**Fig. 1J, Supplemental Fig. 2D**). Surprisingly, +CV-down-regulated groups of gene’s GO terms were clustered and similar to +HFM-up-regulated groups for terms like metabolism of lipids and steroid metabolic process (GO term genes include: *Cyp27a1, Acaa2, Acox2, Acsl1, Acaa1a*) (**Fig. 1J**). Conversely, +HFM-down and +CV-up groups shared positive regulation of defense response (*Nfkbiz, Duoxa2, Tnfrsf11a*), suggesting for certain processes, there may be genes regulated by both nutritional and microbial inputs.

To determine if any of the changes in gene expression were dependent on the interaction of colonization and nutritional status, we characterized the +CV (CV/GF and CV+HFM/GF+HFM, **Fig. 1K**) and +HFM (GF+HFM/GF and CV+HFM/CV, **Fig. 1L**) responses separately. We observed that for most genes the magnitude and directionality of changes are consistent and positively correlated in the two +CV comparisons (**Fig. 1K**). Previously characterized microbially responsive and defense response genes like Saa1, Reg3g, and Tnf were commonly upregulated in both +CV comparisons (**Fig. 1K, Supplemental Table S2**) (Davison et al. 2017). Numerous metabolism genes were commonly downregulated by microbes including previously characterized microbially-responsive genes *Angptl4, Nr0b2, Klf15*, and *Ces3b* (**Fig. 1K; Supplemental Table S2**).

The genes that respond to HFM exposure with and without microbes followed a similar positive correlation of RNA log_2_ fold changes (**Fig. 1L**). *Hmgcs2*, encoding the enzyme required for the first and rate-limiting step in generating ketone bodies from lipid (Cheng et al. 2019), was commonly upregulated by HFM. *Angptl4, Creb3l3, Acot1*, and *Dgat2*, all involved in lipid metabolism, were also commonly upregulated in both +HFM comparisons (**Fig. 1L**). GO term analysis in +HFM-up genes identified enrichment for metabolism of lipids, peroxisome, and PPAR signaling (**Supplemental Fig. S2D; Supplemental Table S2**). Genes down regulated by HFM included genes involved in the unfolded protein response (*Ddit3, Atf3*), transcription factors (*Nfil3, Crebl2, Klf6, Klf4, Fosb*), and signaling components (*Fgf15, Fgf11, Fzd7, Jag1, Wnt5a*). Inflammatory components (*Nfkbiz, Nlr9b, Duox2, Duoxa2*) were down regulated in at least one of the +HFM comparisons.

### Interactions of microbial and HFM signals at FAO genes

In +CV and +HFM comparisons, while certain genes were significant in only one of two pairwise comparisons (e.g. *Cd36* downregulated in CV+HFM/GF+HFM but not CV/GF; Nfil3 downregulated in CV+HFM/CV but not GF+HFM/GF) only anoikis factor *Bmf* (Hausmann et al. 2011) was significantly expressed in opposite directions (Down in CV+HFM/GF+HFM and up in CV/GF) (**Fig. 1K,L** and **Supplemental Fig. 2A**). To test more specifically for interaction between microbes and HFM, we performed Likelihood Ratio Test (LRT) analysis on our RNA-seq data set across all conditions (**Supplemental Fig. S3**) (Love et al. 2014). While only a limited number of genes (*Rnf144b, Slc25a25, Grpc5a, Bmf*) passed a p-adj threshold of .05 for the LRT analysis, we proceeded to lower this cutoff to identify 621 genes that showed the most potential for interaction (**Supplemental Fig. S3A,B**). Typically, putative interaction genes showed substantial increased or decreased relative expression in one condition (**Supplemental Fig. S3A-D,G-I**). For example, genes that were generally reduced primarily in CV+HFM included *Slc10a1, Rnf144b, Bmf,* and *Crot* (**Fig. 1M**). *Ets2, Slc25a25*, and *Gprc5a* were preferentially elevated in CV+HFM (**Fig. 1N**). In GF+HFM, there was a reduction in *Creb1, Acsl4*, and the lipid droplet promoting *Ascl3* (**Fig. 1O**). This was in contrast to the upregulation of FAO genes *Acsl1, Acox2, Amcar*, and *Acad12* in GF+HFM (**Fig. 1P**). Collectively, this suggests that the FAO pathway is reduced by microbes and induced by HFM.

### The same regulatory regions are capable of integrating microbial and nutritional signals

H3K27ac modification of nucleosomes flanking accessible regulatory regions is associated with activation of transcription of neighboring genes. Analysis of these sites in multiple conditions allows for robust interpretation of the genes, transcription factors and pathways involved in coordinated transcriptional regulation (Shen et al. 2012; Davison et al. 2017; Lickwar et al. 2017). We identified H3K27ac levels genomewide in jejunal IECs in the 4 conditions, expanding our understanding of microbially-responsive and identifying HFM-responsive intestinal regulatory regions genomewide. Like we observed in our RNA-seq (**Fig. 1D**), H3K27ac ChIP-seq replicates separate into the 4 conditions using PCA (**Fig. 2A**). An increased number of replicates for CV+HFM and GF+HFM relative to CV and GF improved the ability to identify differential H3K27ac sites, especially for the CV+HFM/GF+HFM comparison, but also led to a much higher number of identified significant sites relative to other comparisons (**Fig. 2C,D**; **Supplemental Table S3**). Similar to our RNA-seq, the *Angptl4* locus showed a region with both microbially reduced and HFM induced H3K27ac levels, with GF+HFM having the highest relative level of H3K27ac (**Fig. 2B**). While many sites only significantly changed in either +CV or +HFM comparisons, 1255 regulatory regions integrate signals from both microbial and HFM conditions (**Fig. 2C; Supplemental Fig. S4A**). Similar to RNA-seq, we found shared enriched GO terms across +CV and +HFM comparisons. For example, +HFM-up and +CV-down sites were nearest to genes related to lipid metabolism, and response to other organisms was found in +HFM-down and +CV-up groups (**Figs. 1J,2E; Supplemental Fig. S4D**). Diverse patterns of H3K27ac enrichment were also found at promoter, intra- and intergenic regions (**Supplemental Fig. S4B,C**), and H3K27ac sites in all comparisons were associated with significant directional changes in RNA levels with neighboring genes (**Fig. 2F; Supplemental Fig. 4C**). At many loci, including *Acsl1, Mdh1, Fads2*, and *Myc*, we can identify the same relative patterns across conditions for RNA expression levels and individual H3K27ac sites, strengthening the likelihood that we have identified regulatory sites that contribute to the integration of combinations of microbial and nutritional stimuli to alter transcription of neighboring genes (**Fig. 2G**).

### Sites with increased H3K27ac following HFM behave differently depending on the presence of microbiota

Having established the utility of our H3K27ac data, we proceeded to compare significant log_2_ fold change levels for H3K27ac sites differential for the two +CV and two +HFM comparisons (**Fig. 2H,I**). Like RNA-seq (**Fig. 1K**), we observed a general positive correlation for relative H3K27ac log_2_ fold change levels in the two +CV comparisons (**Fig. 2H**). +CV sites with increased H3K27ac enrichment were linked to known microbial response genes like *Reg3g, Reg3B, Saa1, Duox2*, and were similarly regulated transcriptionally (Davison et al. 2017). The regulatory regions in both +CV-down H3K27ac comparisons neighbored lipid metabolism genes. These included FAO components like *Acaa2, Ppara, Acsl5*, and *Slc27a4*, and transcription factors *Klf15, Nr1h4, Zbtb16*, and *Id2* (El Aidy et al. 2013; Camp et al. 2014; Davison et al. 2017). However, unlike our RNA-seq analyses, we identified a surprising separation of sites that show increased H3K27ac enrichment following HFM, with a very small number of sites that are significant in both GF+HFM/GF and CV+HFM/CV comparisons (**Fig. 2I**). Interestingly, no such separation occurs for sites that show reduced enrichment in +HFM or at up and down +CV sites (**Fig. 2H,I**). GREAT GO term analysis for the limited shared +HFM-up sites revealed enrichment of lipid metabolism genes including *Acaa2, Pex11a, Angptl4, Abhd6, Cox6c, Entpd7*, and *Trib1*. The GF+HFM/GF-up only H3K27ac sites showed that lipid metabolism genes including *Acsl1, Acaa1b, Pck1, Apoa1, Lipe*, and *Crot* were also the most significantly enriched (**Fig. 2I**). CV+HFM/CV-up H3K27ac GO terms also included lipid metabolism related genes *Apoa1, Cpt1a, Cpt2, Ppard, Dgat2, Fads1, Fads2, Fasn*, and *Lipg*, many of which are associated with lipid anabolism (Chen et al. 2020) (Fig. 2I). Though not represented by a particular GO Term, we also noted many CV+HFM/CV-up regulatory regions neighbor genes with an established role in intestinal proliferation and ISC identity including *Myc, Notch1, Ephb4, Ephb2, Ccnd1, Acsl2*, and *Sox9* (**Fig. 2I**). These genes were also never present in the GF+HFM/GF-up significant group. Importantly, this suggests these functional processes, including divergent lipid metabolic processes and impacts on proliferation and ISCs, are engaged differentially in response to HFM depending upon microbiota colonization.

To identify sites that had the most substantial interaction between the +CV and +HFM variables, like RNA-seq, we used LRT with a lenient statistical cutoff to identify 3430 interacting H3K27ac sites (**Supplemental Fig. S5; Supplemental Table S4**). Comparison between H3K27ac interaction site linked genes that also had transcriptional interactions revealed 89.1% (197/221) were in agreement (**Supplemental Fig. S5F,G**). The loci that showed agreement included genes and sites high in CV+HFM like *Ets2, Fkbp5*, and *Hk2* and those high in GF+HFM like *Acsl1, Bmf*, and *Edn1* (**Supplemental Fig. S5F,K**). This suggests that aspects of the deviant response to HFM with and without microbes is present in both RNA and H3K27ac levels, and that we have mapped many regulatory regions critical to integrating +CV and +HFM signals **(Fig. 2I; Supplemental Fig. S5**).

### Differential integration of both microbial and nutritional signals at the same PPARA/FAO and ISC regulatory regions

To further explore the relationship between these responsive regulatory regions we conducted overlap analysis of significantly differential H3K27ac sites in each comparison. Rather than preferentially overlap with other +HFM sites, GF+HFM/GF-up sites overlapped with CV+HFM/GF+HFM-down sites while CV+HFM/CV-up sites overlapped with CV+HFM/GF+HFM-up sites (**Fig. 3A**). In fact, almost half (673/1375; 48.9%) of the +HFM-up H3K27ac sites were also microbially responsive (**Fig. 3B**). To clarify which processes were being differentially utilized in the +HFM-up comparisons, we partitioned these H3K27ac sites into two groups: “+HFM-up and +CV-up” (red) which typically had highest enrichment in CV+HFM and “+HFM-up and +CV-down” (blue) which typically had highest enrichment in GF+HFM (**Fig. 3A,B**). Based on our initial identification of proliferation and ISC genes neighboring CV+HFM/CV-up sites (**Fig. 2I**), we asked if the separation in the +HFM H3K27ac response with and without microbes was indicative of transcriptional or chromatin differences along the crypt-villus and proliferation-differentiation axis. First, using a dataset that performed H3K27ac and RNA-seq on separate ISC and enterocyte populations in the adult mouse small intestine (Kazakevych et al. 2017), we found that the +HFM-up and +CV-up H3K27ac were rarely accessible solely in enterocytes and instead were frequently enriched only in ISCs, whereas the +HFM-up and +CV-down group frequently contained H3K27ac sites that were enriched in enterocytes (**Fig. 3C; Supplemental Fig. S6A,B**). +HFM-up and +CV-up (red) sites were enriched in processes including regulation of immune system processes (*Myc, Mmp14, Pglyrp1*, and *Stat3*), regulation of epithelial cell differentiation (*Arntl, Mycn, Pax6*, and *Dmbt1*) and Notch signaling (*Hes7, Notch1, Notch2*, and *Sox9*) (**Fig. 3D,E**). We saw limited enrichment for terms involved in metabolism with the notable exceptions of the glycolysis enzyme *Hk2* and limited FAO components *Cpt1a, Cpt2, Ppard*, and *Prdm16*. *Ppard* and *Prdm16* have recently been characterized to regulate FAO in ISCs (Mihaylova et al. 2018; Stine et al. 2019). Several metabolic genes leading to lipid synthesis *Fads1, Fads2, Fasn, Dgat2*, and *Acacb* were only present in this +HFM-up +CV-up group, with Acacb thought to inhibit FAO in favor of lipid anabolism (**Fig. 3E**) (Abu-Elheiga et al. 2001; Abu-Elheiga et al. 2003). Collectively, these sites and their associated genes are indicative of a surplus of energy, lipogenesis, and proliferation in CV+HFM (**Fig. 3E**).

**Figure 3:**
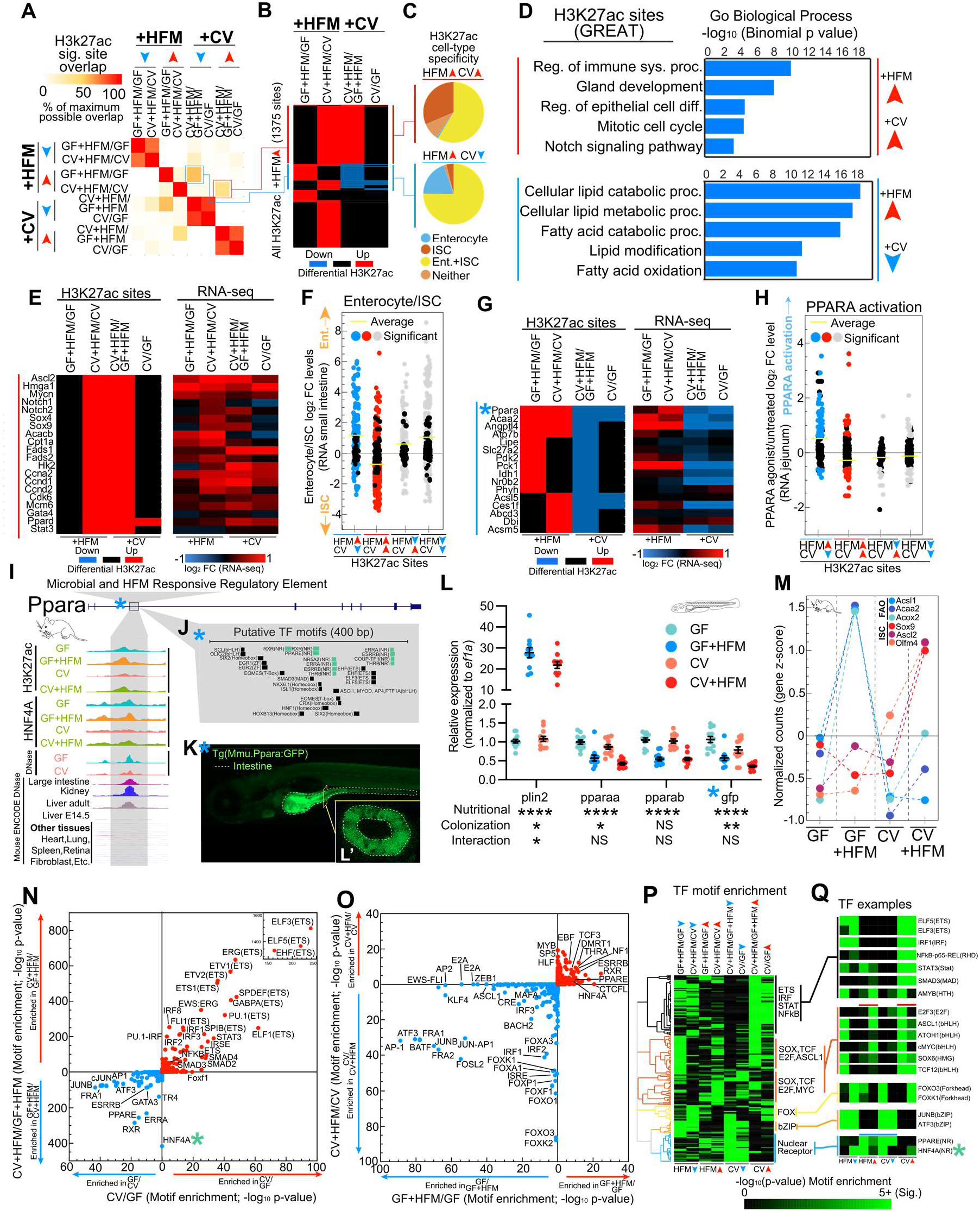
The same regulatory regions can integrate microbial and nutritional signals. **(A)** Pairwise comparison of maximum overlap of coincident significant H3K27ac sites for 8 directional significance groups identifies microbially responsive regulatory regions with CV+HFM/CV-up also being CV+HFM/GF+HFM-up (red) and GF+HFM/GF-up also being CV+HFM/GF+HFM-down (blue). **(B)** Heatmap of differential comparisons for all +HFM-up sites shows the proportion that is also microbially responsive. **(C)** Pie charts for red and blue +HFM-up groups that show the proportion that overlap with enterocyte and ISC regulatory regions(Kazakevych et al. 2017). **(D)** GREAT GO term enrichment for red and blue +HFM-up H3K27ac groups. **(E)** Heatmap of example red +HFM-up and +CV-up H3K27ac sites and their linked gene’s RNA-seq log_2_ fold change, including many loci associated with ISCs and proliferation. **(F)** Different combinations of differential H3K27ac sites that are both +HFM and +CV responsive show that only red sites that are +HFM-up and +CV-up are linked to genes that are preferentially expressed in ISCs relative to enterocytes. **(G)** Heatmap of example blue +HFM-up H3K27ac sites and their linked gene’s RNA-seq log_2_ fold change. Blue asterisk marks Ppara regulatory region that is characterized in I-L. **(H)** Different combinations of differential H3K27ac sites that are both +HFM and +CV responsive show that only blue sites that are +HFM-up and +CV-down are linked to genes that are activated by PPARA. **(I)** Exploded view of H3K27ac region at Ppara locus from **(G)** that is +HFM-up and +CV-down showing signal across conditions for H3K27ac, HNF4A, and accessible chromatin for jejunum, and numerous other tissues. **(J)** Putative TF motifs at the *Ppara* regulatory region include multiple nuclear receptor sites, including a PPARE. **(K)** Transgenic *Tg*(*Mmu.Ppara:GFP*) F1 zebrafish with mouse *Ppara* regulatory region upstream of the cFOS minimal promoter driving GFP shows expression largely limited to the anterior intestine of zebrafish. K’ boxed inset shows a confocal cross section confirming the signal is specific to IECs. **(L)** qRT-PCR of a four condition experiment using whole *Tg*(*Mmu.Ppara:GFP*) zebrafish shows similar responses to colonization and high fat meal for *pparaa* and *gfp*. Significance calls for colonization, nutritional and interaction based on 2-factor ANOVA. *p-value=<.05, **p-value=<.01, and ****p-value =<.0001. **(M)** FAO and ISC associated genes showing particular expression in GF+HFM and C+HFM, respectively (**Supplemental Table S2**). **(N)** Motif enrichment at DNase sites linked to +CV H3k27ac significance groups comparing CV/GF and CV+HFM/GF+HFM sites. For each significance +CV group the reciprocal direction is used as the background (e.g. CV/GF-up input versus CV/GF-down background). –log_10_ p-values are plotted. Both directions are plotted on the same axis with each analysis separated by colored arrows. HNF4A motif (green asterisk) was not differentially enriched between CV/GF directional H3K27ac sites, but was substantially enriched in CV+HFM/GF+HFM-down sites relative to CV+HFM/GF+HFM-up sites. **(O)** Same as N for GF+HFM/GF versus CV+HFM/CV. **(P)** Clustering of enrichment motif score (−log_10_ p-value) for TF motifs that are present in multiple comparisons for 8 directional significance groups. Data is shared with N and O. **(Q)** Example TF motif patterns including sites coincident with red (+HFM-up and +CV-up) and blue (+HFM-up and +CV-down) H3K27ac sites.

To determine if these +HFM-up and +CV-up (red) sites were preferentially associated with ISC versus enterocyte expressed genes, we looked at different combinations of H3K27ac sites that responded to both microbial and nutritional signals based on the expression of their neighboring genes in enterocytes/ISCs **(Fig. 3F**) (Kazakevych et al. 2017). The +HFM-up and +CV-up (red) sites were linked to genes expressed preferentially in ISCs relative to enterocytes (**Fig. 3F**). Importantly, including +HFM-up and +CV-down (blue), no other sites that responded to both +HFM and +CV signals were associated with ISCs and enterocyte progenitor genes (**Fig. 3F; Supplemental Fig. S6C-E**). To confirm this observation, we leveraged a dataset identifying genes differentially expressed in 5 zones along the crypt-villus axis in the mouse small intestine (Moor et al. 2018). Comparing those patterns to our H3K27ac data, we observed substantial preference for increased enrichment of crypt associated H3K27ac sites in CV+HFM/GF+HFM and decreased enrichment was associated with the villus (**Supplemental Fig. S6E**). This suggests that these red ISC sites are initiated primarily in both the presence of +HFM and +CV signals, notably the CV+HFM condition, and are less activated by HFM in the absence of microbes (**Fig. 3F**).

In contrast, genes neighboring H3K27ac sites that were +HFM-up and +CV-down (blue) (**Fig. 3B**) were enriched for lipid metabolism GO terms like cellular lipid catabolic process and FAO (**Fig. 3D**). The GO terms included genes *Angptl4, Acaa2, Acsl5*, and the gluconeogenic regulator *Pck1*, and these genes typically also showed similar expression patterns at the level of RNA-seq and H3K27ac (**Fig. 3G**). In addition, a regulatory region at the FAO regulator, *Ppara*, was in the +HFM-up and +CV-down group (**Fig. 3G**). FAO genes and other genes associated with energy production are transcriptionally activated by the lipid-liganded PPARA to facilitate the production of ATP from lipids (Dubois et al. 2017). To identify if +HFM-up and +CV-down (blue) sites were associated with PPARA signaling, we queried a published dataset of genomewide gene expression in the mouse jejunum following activation of PPARA by the agonist WY14643 in both WT and *Ppara* knockout mice (Bunger et al. 2007). +HFM-up and +CV-down H3K27ac sites were substantially linked to PPARA target genes (**Fig. 3H; Supplemental Fig. S6D,F**). Importantly, no other groups of sites, including +HFM-up and +CV-up (red), showed a clear response to PPARA activation on average. This suggests that only sites that integrated the particular combination of both +HFM-up and +CV-down signals were PPARA targets (**Fig. 3H**). Consistent with this finding, comparing expression levels after PPARA activation versus HFM response or microbial response revealed a positive and negative correlation, respectively (**Supplemental Fig. S6G-J**).

### Mouse *Ppara* enhancer shows evidence of conserved intestinal expression and microbial and HFM responsiveness in transgenic zebrafish

We next sought to characterize the *Ppara* regulatory region that may integrate +HFM and +CV signals (**Fig. 3G,I; Supplemental Fig. S6K**). The H3K27ac enhancer we identified at mouse *Ppara* is bound by HNF4A, displays accessibility largely restricted to adult digestive tissues, and contains a PPARE and other nuclear receptor binding sites (**Fig. 3IJ**). To test if that enhancer was capable of integrating microbial and nutritional signals in IECs, we cloned it into a zebrafish GFP reporter assay. Stable transgenic zebrafish harboring the *Ppara* mouse enhancer reporter drove expression largely in IECs (**Fig. 3K**), indicating that it mediates IEC expression in zebrafish as well as mouse. We found that, as in mouse, both zebrafish *pparaa* and GFP were down regulated following colonization (**Fig. 3L**). However, the GFP reporter or zebrafish *pparaa* was not activated by HFM after 6h in zebrafish (Semova et al. 2012; Zeituni et al. 2016). Instead, both were concordantly reduced by HFM, suggesting that this mouse regulatory region and *pparaa* in zebrafish still may integrate microbial and nutritional signals in a conserved manner in IECs (**Fig. 3L**). Collectively, this suggests that HFM preferentially influences H3K27ac sites around genes involved in ISCs and proliferation with microbes and PPARA signaling and FAO without microbes. This effect can also be seen at key FAO markers and ISC markers like *Olfm4* and *Sox9* at the level of RNA in mouse (**Fig. 3M**).

### HNF4A motif is differentially enriched in microbially suppressed H3K27ac sites only in the presence of HFM

We next utilized transcription factor motif enrichment to identify those enriched at regulatory regions with differential H3K27ac utilization in the 4 conditions (**Fig. 3N-Q**). Common +CV-up motifs include established microbially-responsive TFs like NF-κB, IRF, and STAT (**Fig. 3N**). We previously showed HNF4A motifs are not significantly differentially present at CV/GF sites that show increased or decreased H3K27ac enrichment (**Fig. 3N**) (Davison et al. 2017). Surprisingly, in the presence of HFM, the HNF4A motif was enriched at sites that have reduced H3K27ac with a microbiota (CV+HFM/GF+HFM-down). We next clustered significant motifs for each comparison to identify patterns of their enrichment in +CV and +HFM conditions (**Fig. 3P,Q**). +HFM-up and +CV-up (red) conditions included several motifs associated with proliferation including E2f3, Ascl1, and cMYC. Both PPAR and HNF4A showed motif enrichment in +HFM-up and +CV-down comparisons suggesting they are also involved in integrating microbial and HFM signals associated with the blue H3K27ac sites group (**Fig. 3N-Q; Supplemental Fig. 6L,M**).

### The association of *Hnf4a* with the response to microbes is due to its impact on enterocyte differentiation and the crypt-villus axis

To understand how HNF4A binding contributes to transcriptional integration of microbial and nutritional signals, we performed HNF4A ChIP-seq in the four conditions. HNF4A occupancy is reduced at many binding sites in the CV conditions suggesting HNF4A is more active in GF and this activity is suppressed by microbes (Davison et al. 2017). PCA analysis shows that unlike RNA-seq and H3K27ac, HNF4A does not separate substantially by all four conditions, with only CV datasets separating along axis 1 (**Fig. 4A**). Consistent with this, we saw that HNF4A binding recovered occupancy following HFM in the CV+HFM condition **(Fig. 4B,C; Supplemental Table S5**). Differential HNF4A binding analysis identified distinct changes, with +HFM generally increasing HNF4A occupancy and +CV generally decreasing HNF4A occupancy (**Fig. 4C-E; Supplemental Fig. 7A-D**). While provocative, this behavior appeared to be consistent at most sites across the genome, thereby limiting our ability to interpret meaningful biological differences in HNF4A occupancy levels from individual sites (**Fig. 4C; Supplemental Fig. 7E-L**).

**Figure 4:**
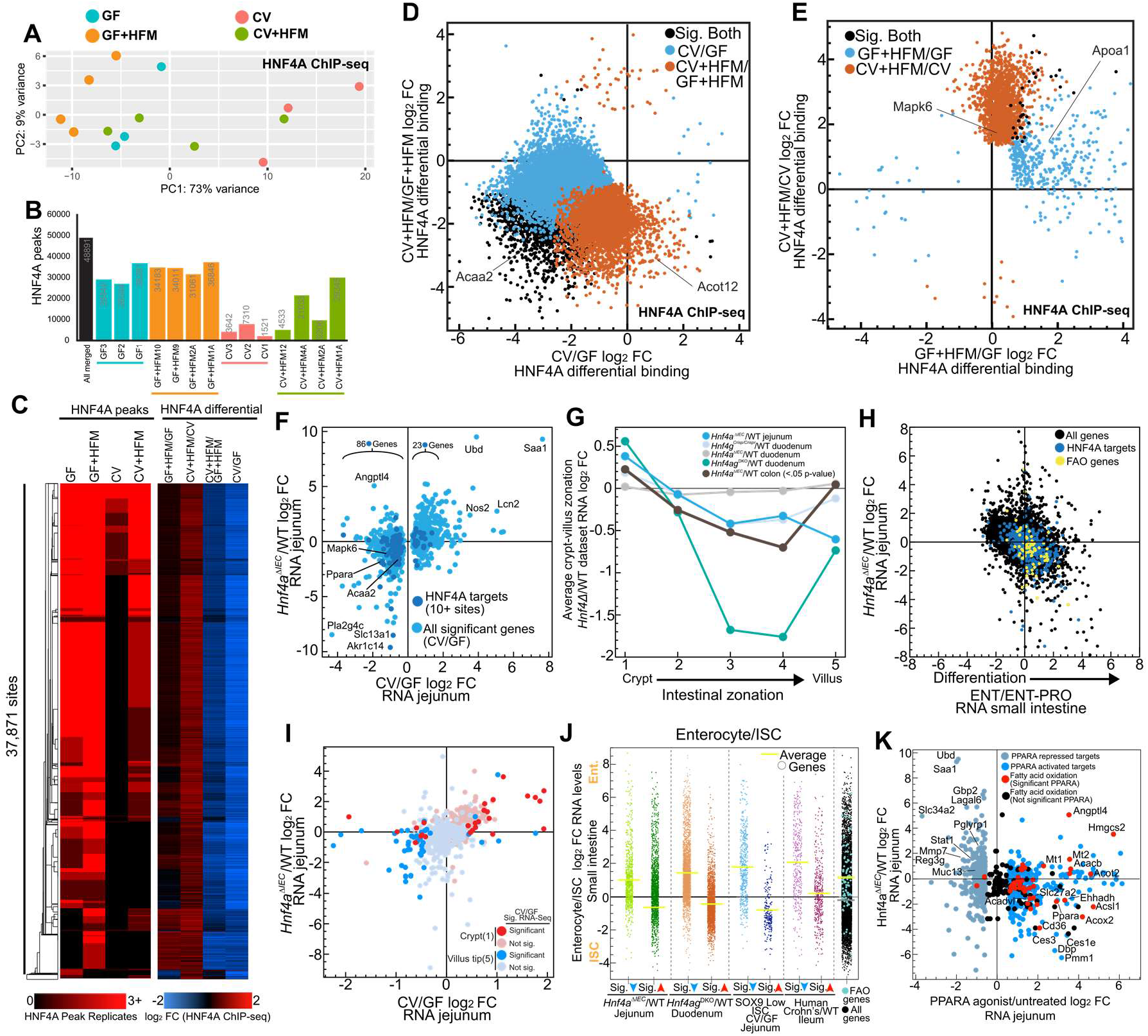
HNF4A’s role in promoting IEC differentiation explains correlation between *Hnf4a* loss and microbially responsive genes. **(A)** PCA of HNF4A ChIP-seq normalized counts shows the CV condition deviates from all other conditions. **(B)** HNF4A peak numbers across GF, GF+HFM, CV and CV+HFM for merged HNF4A peaks with overlap from at least two Hnf4a replicates. **(C)** Clustered heatmap of HNF4A binding sites with peaks called in at least 2 replicates compared to log_2_ fold change for each of the 4 comparisons. **(D)** Scatterplot of significantly different log_2_ fold change for CV/GF and CV+HFM/GF+HFM HNF4A occupancy. **(E)** Same as D for GF+HFM/GF and CV+HFM/CV HNF4A occupancy. **(F)** Scatterplot comparison between significantly differential microbial responsive genes and *Hnf4a*^ΔIEC/^WT in the jejunum (Verzi et al. 2013). Genes with more than 10 HNF4A sites (dark blue). **(G)** Comparison of genes that are preferentially expressed in five compartments along the crypt-villus axis for various *Hnf4* deletion mutants in the intestine (Darsigny et al. 2009; Verzi et al. 2013; Moor et al. 2018; Chen et al. 2020). **(H)** Scatterplot comparing small intestine RNA-seq log_2_ fold change for enterocyte/enterocyte progenitor versus *Hnf4a*^ΔIEC/^WT. 10+ HNF4A binding sites/targets (blue) contribute directly and indirectly to FAO genes (yellow) activation preferentially in enterocytes (Verzi et al. 2013; Kim et al. 2014). **(I)** Scatterplot comparing jejunal microbial response (CV/GF) to *Hnf4a*^ΔIEC^/WT in jejunum colored for genes preferentially expressed in the crypt (red) and villus tip (blue) (Verzi et al. 2013; Moor et al. 2018). **(J)** RNA-seq log_2_ fold change for small intestinal enterocyte/ISC RNA levels for groups of significantly differential down (blue arrow) and up (red arrow) genes from numerous published data set showed a common impact of *Hnf4a*^ΔIEC^/WT(Verzi et al. 2013), *Hnf4ag^DKO^*/WT (Chen et al. 2020), CV/GF in sorted ISCs (Peck et al. 2017), and human ileal Crohn’s (Haberman et al. 2014) on the crypt-villus/proliferation-differentiation axis (Kazakevych et al. 2017). FAO genes are also more highly expressed in enterocytes versus ISCs (**Supplemental Table S6**). Yellow bars refer to the average Enterocyte/ISC log_2_ fold change RNA levels for each group. Gray bar represents the average for all genes except FAO genes. **(K)** Scatterplot comparing PPARA-activated genes versus *Hnf4a*^ΔIEC^/WT RNA levels in mouse jejunum identify PPARA targets and FAO genes are commonly reduced in *Hnf4a*^ΔIEC^.

We instead focused on interpreting the relatively stable position of HNF4A binding sites across occupied conditions as well as the relationship between HNF4A-activated and microbially-suppressed genes we have seen in zebrafish (Davison et al. 2017). Using a previously published dataset of mice lacking *Hnf4a* in jejunal IECs (*Hnf4a*^ΔIEC^) (Verzi et al. 2013), we found that HNF4A-activated genes are also more highly expressed in the GF condition relative to CV. That suggests the relationship between the HNF4A regulon and microbially-responsive genes is conserved in mouse jejunum, with a similar relationship also seen in mouse colon (**Fig. 4F, Supplemental Fig. 7M**). We also saw that genes upregulated in *Hnf4a*^ΔIEC^ mice are correlated with those upregulated following colonization (**Supplemental Fig. 7N**), with inflammation related genes like Saa1 being consistently upregulated by *Hnf4a* loss and presence of microbes across species (**Fig. 4F**) (Davison et al. 2017).

To distinguish between direct and indirect targets, we also identified which of these mouse genes are bound by a neighboring HNF4A site. HNF4A binding sites are numerous in the intestine with typically 15,000-30,000+ binding sites in IECs (Verzi et al. 2010; Chahar et al. 2014; Davison et al. 2017; Qin et al. 2018). However, transcriptional level changes in *Hnf4a*^ΔIEC^ mice have been shown to be correlated with not just binary presence or absence, but the cumulative number of neighboring HNF4A binding sites for a particular gene (**Supplemental Fig. 7O-P**) (Verzi et al. 2013; Chahar et al. 2014; Chen et al. 2019b). We found that genes with abundant neighboring HNF4A binding sites (10+ binding sites) are more likely to lose expression in both Hnf4a^ΔIEC^ and following colonization with microbes (**Fig. 4F, Supplemental Fig. 7Q-V**). Conversely, by this metric, genes induced by microbial colonization and in *Hnf4a*^ΔIEC^ are less frequently direct HNF4A targets. Importantly, even at the remaining loci with little or no evidence of being a direct HNF4A target, a correlation between *Hnf4a*^ΔIEC^ and CV/GF expression levels remained for both up- and down-regulated genes (**Fig.4F, Supplemental Fig. 7Q,R**). This implicates both direct, indirect, or non-traditional HNF4A functions as contributors to the correlation between HNF4A- and microbially-regulated genes in the jejunum. This is consistent with findings of HNF4A functioning primarily as an activator that defines the chromatin landscape in IECs (Verzi et al. 2013; San Roman et al. 2015; Chen et al. 2019a), and HNF4A in regulating genes that are reduced in expression following colonization with microbes (Davison et al. 2017). This also suggests that the impact of HNF4A function in response to microbiota and other functions may be cumulative and distributed across multiple regulatory regions for a particular gene (Chen et al. 2021).

### Microbial and nutritional stimuli influence the crypt-villus axis leading to differential utilization of Hnf4a and differentiation programs

Considering the relatively uniform, numerous positions of binding sites of HNF4A in IECs, we investigated if more global changes in cell identity or cellular programs explains the relationship between HNF4A and microbial colonization. HNF4A and HNF4G initially were identified as genome-wide positive regulators of enterocyte differentiation with reduced HNF4A activity in proliferating cells (Babeu et al. 2009; Cattin et al. 2009; Verzi et al. 2010). As a result, loss of *Hnf4a* leads to reduced expression of differentiation genes and increased proliferation genes in an intestinal cell line, and HNF4A and HNF4G are believed to work together to promote brush border genes (Chen et al. 2021). A direct impact for HNF4A and HNF4G in regulating FAO in ISCs has also been suggested (Chen et al. 2020). We find loss of *Hnf4a* resulted in increased expression of crypt genes and a decrease of genes expressed in the differentiating villus in the jejunum, colon, or the simultaneous loss of *Hnf4a* and *Hnf4g* in duodenum (**Fig. 4G**) (Darsigny et al. 2009; San Roman et al. 2015; Chen et al. 2019b). Similarly, comparison of genes that are preferentially expressed in terminally differentiated enterocytes relative to enterocyte progenitors or ISCs identified a negative correlation across all genes with expression in *Hnf4a*^ΔIEC^ mouse jejunum, consistent with HNF4A directly and indirectly promoting enterocyte differentiation (**Fig. 4H; Supplemental Fig. 7X**). To simultaneously interrogate the impact of HNF4A and microbes on the crypt-villus axis, we overlaid the preferential position of a gene’s expression along the crypt-villus axis relative to microbial response and *Hnf4a* dependence (**Fig. 4I**) (Moor et al. 2018). Genes that were both microbially suppressed and lose expression in *Hnf4a*^ΔIEC^ were preferentially expressed in the differentiated villus. In contrast, microbially-activated and genes with increased expression in *Hnf4a*^ΔIEC^ coincidentally were preferentially expressed in the crypt, directly implicating HNF4A and the response to microbes in regulating differences in the crypt-villus axis and enterocyte differentiation (**Fig. 4I**).

In support of the hypothesis that microbial responses impact the underlying differentiation status of IECs, we queried a published RNA-seq dataset of sorted jejunal ISCs in separate GF and CV conditions. We not only observed the previously identified increased expression of proliferation and ISC genes following microbial exposure, but also significantly reduced expression of genes preferentially expressed in enterocytes vs ISCs (**Fig. 4J**) (Peck et al. 2017). This suggests that, even when constrained solely to a population of ISCs, differences in the crypt-villus or ISC-enterocyte differentiation axis can be substantially perturbed by microbes and are unlikely to solely be driven by differences in IEC subtype composition between conditions. Differential expression of genes along the crypt-villus axis may also explain why a relationship between inflammatory phenotypes and HNF4A are seen in human Crohn’s disease (**Fig. 4J**) (Haberman et al. 2014; Davison et al. 2017).

We speculated that HNF4A may promote preferential expression of FAO genes in differentiated enterocytes in the GF and GF+HFM conditions relative to CV+HFM. We found that FAO genes are preferentially expressed in enterocytes versus ISCs and enterocyte versus enterocyte progenitors in normal chow-fed conditions (**Fig. 4H-J, Supplemental Fig. 7X; Supplemental Table S6**). Similarly, loss of *Hnf4a* in jejunum results in loss of expression of FAO genes, PPARA targets generally, and *Ppara* itself (**Fig. 4K**) (Bunger et al. 2007; San Roman et al. 2015; Davison et al. 2017). This suggests that some of the capacity for FAO in the jejunum relies on HNF4A potentiating an enterocyte differentiation program upstream of *Ppara* and FAO targets (Chen et al. 2020).

### Condition specific H3K27ac changes are restricted to enterocyte- and ISC-specific regulatory regions and impacted by Hnf4a binding

Since 2h after a HFM is likely too soon to induce major changes in IEC cell-type abundance (Cheng and Leblond 1974; Al-Dewachi et al. 1975), the enrichment of HNF4A motifs in GF+HFM (**Fig. 4N-Q**) and differential utilization of enterocyte and ISC regulatory regions in +HFM-up conditions (**Fig. 3C**) may be due to a global change in the utilization of the enterocyte-differentiation transcriptional program within the larger IEC population (**Fig. 3J**). We generated a limited intestinal genomics database using a previously published dataset of accessible chromatin mapping of ISCs, proliferating transit-amplifying (TA) cells, enteroendocrine cells (EECs), and enterocytes using the Sox9 sorting strategy from adult mouse intestine as a foundation (Raab et al. 2020). Organizing accessible sites merged from these IEC subtype populations with our DNase Hypersensitivity (DHS) data from jejunum identified relatively simple patterns of chromatin accessibility across IEC subtypes (**Fig. 5A**) (Davison et al. 2017). To test the impact of the crypt-villus, ISC-enterocyte differentiation axis, we grouped enterocyte-accessible regions that were also accessible across all other IEC subtype populations (Pan-accessible, includes ISC), sites containing enterocyte accessibility and at least one other cell subtype (Enterocyte-restricted), and enterocyte-specific accessible sites (Enterocyte-specific) (**Fig. 5A**). We similarly grouped ISC-specific and ISC-restricted sites (**Fig. 5A**). Using DHS accessibility from multiple cell types and our published data from colon and ileum, we identified that Pan-accessible sites are frequently accessible constitutively across all cell types while sites with accessibility in fewer types of IECs are less often accessible in other tissues (**Fig. 5A**) (Mouse et al. 2012; Camp et al. 2014).

**Figure 5:**
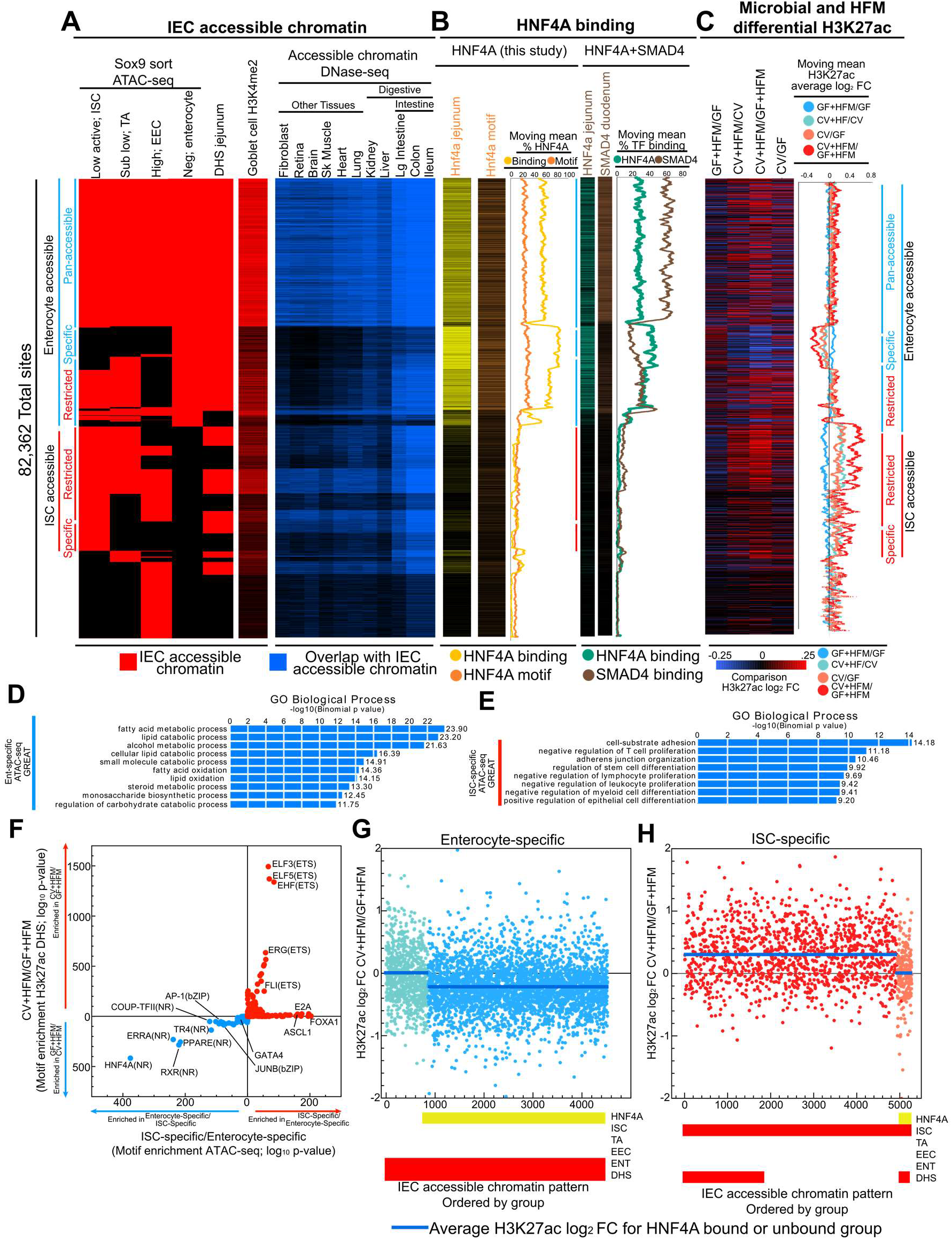
HNF4A-dominated enterocyte and HNF4A-absent ISC regulatory regions behave distinctly in response to microbes and high fat meal. **(A)** Organized heatmap of merged accessible chromatin peak calls for sites that were significantly enriched in at least one of the four sorted IEC cell types (red) (Raab et al. 2020). Jejunal DHS sites (Davison et al. 2017) and H3K4me2 sites from goblet cells (Kim et al. 2014) are also included. Groups of sites that are accessible in enterocytes and ISCs are further broken down into whether they are accessible in other (restricted) or only their IEC type (specific). A subset of sites are accessible across IEC subtypes (pan-accessible). Corresponding accessible chromatin data overlap with other mouse tissues (blue) identifies patterns of enrichment (Mouse et al. 2012; Camp et al. 2014). Groups of sites that correspond to enterocyte sites are marked with a straight blue line and ISC but not enterocytes sites are marked with straight red lines. **(B)** Heatmap for HNF4A binding (yellow) and computationally detected HNF4A motif (orange) for sites ordered as in A. This data is summarized with descending moving means (500 site window, 1 site step). A similar pattern is seen using a previously published HNF4A ChIP-seq (teal) (Verzi et al. 2013). Duodenum SMAD4 (brown) (Chen et al. 2019b). **(C)** Heatmap for H3K27ac differential binding for sites ordered as in A. These data are summarized with descending moving means (500 site window, 1 site step). **(D)** GREAT GO term enrichment for enterocyte-specific sites. **(E)** GREAT GO term enrichment for ISC-specific sites. **(F)** Motif enrichment at ATAC-seq sites linked to ISC-specific and Ent-specific groups versus CV+HFM and GF+HFM groups. For each group the reciprocal direction is used as the background (i.e. ISC-specific input versus Ent-specific background). –log_10_ p-values are plotted. Both directions are plotted on the same axis with each analysis separated by colored arrows. **(G)** Groups of accessible sites (red bars) further organized by if they are bound by HNF4A (yellow bars) show dependence on HNF4A binding for decreased enrichment in CV+HFM/GF+HFM H3K27ac signal at enterocyte-specific sites and increased **(H)** ISC-specific sites on average.

We overlapped these accessible chromatin sites with HNF4A binding from our data and identified that up to 60% of the Pan-accessible sites were also bound by HNF4A in IECs (**Fig. 5B**). Enterocyte-specific sites rose to a substantial 80% overlap with HNF4A binding (**Fig. 5B**). In contrast, sites without enterocyte accessibility, including the large group of ISC-specific and -restricted sites, had limited to no HNF4A binding (~10%) in our IEC preparation (**Fig. 5B**). We confirmed this overall pattern using a previously published data set for HNF4A binding in mouse jejunal IECs (**Fig. 5B**) (San Roman et al. 2015; Chen et al. 2019b). We also computationally scored each site for the presence of an HNF4A motif, which would be independent of influence from chromatin utilization or binding in a particular cell type. We detected the HNF4A motif at 30% of the sites with enterocyte accessibility and only 10% of the sites without accessibility in enterocytes. This suggests that sites that are accessible in ISC, but not enterocytes, are not typically conventionally bound by HNF4A in any circumstance. Importantly, though *Hnf4a* is expressed in ISCs, our analysis indicates that identification of HNF4A motif enrichment in ISC accessible chromatin is likely due to sites coincidently accessible in enterocytes (**Fig. 5B**) (Chen et al. 2020). SMAD4 has recently been shown to contribute to enterocyte differentiation with *Hnf4a* (Chen et al. 2019b). While we see substantial coincidence of SMAD4 and HNF4A binding, the proportion of sites that are bound by SMAD4 at enterocyte-specific sites is reduced (~20%) as compared to pan-enterocyte sites (60%), suggesting HNF4A contributes more exclusively to this subset of enterocyte-specific accessible sites (**Fig. 5B**).

Finally, to query the impact of microbial and nutritional signals on the different types of regulatory regions, we compared the relative H3K27ac signal for each site for our 4 comparisons (**Fig. 5C**). For pan-accessible and enterocyte-restricted sites we saw relatively no change in H3K27ac on average. However, at enterocyte-specific sites we identify consistent H3K27ac changes based on the +HFM or +CV comparison (**Fig. 5C**). CV+HFM/CV and CV/GF show reduction of H3K27ac signal at these enterocyte-specific sites. This reduction is even more substantial in CV+HFM/GF+HFM, with the GF+HFM/GF comparison increasing. This suggests that generally microbes, especially in combination with HFM, reduce the utilization of enterocyte-specific sites. We identified the opposite effect when looking at ISC-specific sites which showed substantial H3K27ac signal increases in CV/GF, CV+HFM/CV, even greater increases when comparing CV+HFM/GF+HFM and reduced signal at GF+HFM/GF (**Fig. 5C**). This difference suggests a major impact of HFM, in the presence or absence of microbes, is manipulation of the related crypt-villus, ISC-enterocyte, or proliferation-differentiation axis (**Supplemental Fig. 5E**). We found that enterocyte-specific sites are enriched near genes involved in carbohydrate metabolism, FAO, and lipid catabolism including *Abcd3, Acaa2, Acox1, Crot, Ppargc1a, Acsl1*, and *Ppara*. In contrast, ISC-specific sites are near genes involved in stem cell differentiation (**Fig. 5D,E**). HNF4A and other NR motifs that were enriched in GF+HFM amplified regulatory regions are also enriched in these enterocyte-specific sites (**Fig. 5F**). Though this property may be attributable to enterocyte-specific regions behaving as a group or reminiscent of a change in the proportional number of a given cell-type, we surprisingly found that only enterocyte-specific sites also bound by HNF4A showed altered H3K27ac changes on average (**Fig. 5D-F**). This suggests that HNF4A binding influences interacting microbial and nutritional enterocyte-specific regulatory regions.

## Discussion

Physiologic responses by the intestine to microbiota and diet must be understood as dynamic processes. The postprandial response typically takes 8 hours for lipid absorption (Uchida et al. 2012) and can be influenced by host history, including diet, microbes, and circadian rhythm (Uchida et al. 2012; Mukherji et al. 2013; Wang et al. 2017; Martinez-Guryn et al. 2018). Colonization of GF mice has leads to transcriptional responses that remain dynamic for up to 30 days and are dependent on the composition of the microbial community (El Aidy et al. 2012; El Aidy et al. 2013). Simultaneously, the intestinal epithelial layer turns over every 3-5 days with the potential to alter IEC subtypes and crypt-villus architecture while simultaneously making acute transcriptional changes on the order of minutes (Cheng and Leblond 1974; Al-Dewachi et al. 1975; El Aidy et al. 2013; Mah et al. 2014; Nyima et al. 2016; Shamir et al. 2016; Zeituni et al. 2016; Haber et al. 2017; Mucunguzi et al. 2017; Xie et al. 2020; Heppert et al. 2021). We aimed here to understand how the intestine integrates two distinct conditions simultaneously, and specifically how microbes modify the response to a high fat meal. Focusing on a single early stage in the postprandial process, we identified a response with substantial complexity but also with clear contributions from PPARA, proliferation pathways, and HNF4A-driven enterocyte differentiation.

At the level of H3K27ac and RNA-seq, our results indicate that PPARA signaling is preferentially engaged following HFM in the absence of microbes. Nuclear receptor and PPARA signaling has previously been associated with gene expression in GF that is suppressed by microbes (Backhed et al. 2004; El Aidy et al. 2013; Mukherji et al. 2013); Camp et al. (2014); (Martinez-Guryn et al. 2018; Tazi et al. 2018; Araujo et al. 2020). The egg yolk meal used here is also rich in fatty acids, which could act as PPARA ligands (Wood et al. 2021). PPARA and CREB1 regulatory networks have been associated with the acute transcriptional response to high and low fat meals, respectively (Nyima et al. 2016; Mucunguzi et al. 2017). Combined, this may explain why GF+HFM condition had the highest relative expression level of genes involved in PPARA signaling (**Figs. 1P, 3H**). We speculate that the high level of *Ppara* and FAO gene transcription in GF and GF+HFM may contribute to reduced accumulation and absorption of dietary lipid as compared to CV following HFM (Semova et al. 2012; Martinez-Guryn et al. 2018; Tazi et al. 2018; Araujo et al. 2020). FAO may also be influenced by microbial metabolites such as acetate through the AMPK/PGC-1α/PPARα signaling pathway or by c-Jun/NCoR regulation (Backhed et al. 2007; Mukherji et al. 2013; Araujo et al. 2020). We found the increased expression of PPARA signaling genes in GF and GF+HFM was linked to preferentially utilized H3K27ac regulatory regions in these conditions (**Fig. 3H**). This same subset of regulatory regions also responded with decreased H3K27ac to microbes, suggesting aspects of the microbial and HFM response are integrated at the same sites and mediated by PPARA signaling.

Interestingly, we find that the same PPARA-associated genes that are induced by HFM are also induced by fasting (**Supplemental Fig. F5I,J**). Fasting liberates stored fat to produce PPAR-activating fatty acids that can also be burned by FAO to produce ATP (Kersten et al. 1999). GF mice may therefore be primed to initiate FAO to utilize a HFM as compared to CV mice, potentially due to enterocyte abundance, other IEC identity preferences, or other forms of regulation (Mukherji et al. 2013; Araujo et al. 2020), and elevated PPARA levels may more readily be activated by an influx of lipids in the GF state (**Supplemental Fig. S8**). Furthermore, activation of PPARA reduces lipid droplet levels specifically in jejunum (Karimian Azari et al. 2013), potentially with regulatory contributions from LXR (Colin et al. 2008). We were also interested to see that pharmacological activation of PPARA not only led to activation of FAO genes that are typically suppressed by microbiota, but also suppressed genes normally induced by microbiota such as Reg3g and Saa1 (**Supplemental Fig. 6H**) (Bunger et al. 2007). This raises the interesting possibility that PPARA signaling directly opposes proinflammatory programs in the small intestine (Mukherji et al. 2013).

In contrast to PPARA signaling, ISC and proliferation components were preferentially engaged by HFM in the presence of microbes. Increased IEC proliferation has been associated with colonization with a microbiota in both mouse and zebrafish (Rawls et al. 2004; Cheesman et al. 2011; El Aidy et al. 2013; Park et al. 2016; Peck et al. 2017). Chronic high-fat diet is also associated with changes in crypt size and increased intestinal proliferation (Mao et al. 2013; Mah et al. 2014; Beyaz et al. 2016; Xie et al. 2020). In Drosophila, only in the presence of microbes does a high-fat diet induce proliferation in the intestine (von Frieling et al. 2020). We found that regulatory regions that have increased H3K27ac signal in response to both microbes and HFM (+HFM-up +CV-up) are associated with ISC-specific sites, and neighboring genes are preferentially expressed in ISCs versus enterocytes (**Fig. 3C,F**). This suggests that a major alteration in transcriptional changes associated with the crypt-villus axis is underway as early as 2h after HFM in the presence of microbiota. Based on typical cell proliferation rates, we consider it unlikely that substantial changes in cell type abundance occurred within 2h (Cheng and Leblond 1974; Al-Dewachi et al. 1975). Whereas different cell subtypes typically display distinct transcriptional and chromatin profiles, it is possible that physiological adaptation causes certain cell subtypes to adopt characteristics typically associated with other cell types or metabolic programs. Indeed, we find that GF jejunal ISCs maintain higher expression of genes associated with enterocytes relative to ISCs following 2 weeks of microbial colonization (**Fig. 4J**) (Peck et al. 2017). This suggests that enterocyte-enriched metabolic pathways like FAO communicate with pathways regulating epithelial renewal. Indeed, loss of *Cpt1a* expression in IECs leads to reduced FAO and proliferation (Mihaylova et al. 2018; Mana et al. 2021), and loss of FAO in *Hnf4ag^DKO^* or by chemical suppression is coincident with reduced ISC markers, but increased transit amplifying proliferation markers (Chen et al. 2020). In contrast, activation of FAO with a PPARA agonist results in suppression of the MYC proliferation program in the intestine (Bunger et al. 2007). These findings do not preclude PPAR signaling and FAO components from being important and more highly expressed in enterocytes (Fig. 4J). This implies a complicated relationship between ISCs, proliferation, and FAO/PPARA signaling in IECs that is not yet fully understood in the context of microbial colonization (Mukherji et al. 2013).

HNF4A may sit at the intersection of microbial and nutritional signals and to intestinal adaptation generally (Dubois et al. 2020). Here we identify a conserved correlation between microbially-regulated genes and HNF4A-regulated genes in mouse jejunum (**Fig. 4E-I**) (Davison et al. 2017). We propose that the correlation is driven by the related function of HNF4A promoting enterocyte differentiation relative to proliferation, and microbial colonization promoting proliferation relative to differentiation. HNF4G, which also contributes to enterocyte differentiation, and binds at the same sites as HNF4A in IECs, is transcriptionally microbially-suppressed in the CV+HFM/GF+HFM comparison suggesting it may also contribute to intestinal adaptation (**Supplemental Fig. 6L; Supplemental Table S2**) (Davison et al. 2017; Lindeboom et al. 2018; Chen et al. 2019b; Raab et al. 2020). Understanding contributions of HNF4A to the genomic regions that regulate microbial responsiveness is complicated. HNF4A is proposed to function primarily as an activator genome-wide in IECs, though there are reports of HNF4A repressor function (Rodriguez et al. 1998; Martinez-Jimenez et al. 2010; Santangelo et al. 2011; Qu et al. 2018). While microbial responsive enhancers in CV and GF are enriched for HNF4A binding, GF enhancers are not substantially more often HNF4A bound, despite *Hnf4a* deletion being associated with activation of genes most highly expressed in GF (**Fig. 4F,N-Q**) (Davison et al. 2017). We found a higher number of HNF4A binding sites per gene at microbially-suppressed genes than at microbially activated genes. Interestingly, even at microbially-suppressed genes not bound by HNF4A, the correlation persisted between microbially-regulated and HNF4A dependent genes (**Fig. 4F; Supplemental Fig. S7Q-V**).

In endodermal organs like liver and intestine HNF4A is a known regulator of lipid metabolism (Yin et al. 2011; Chen et al. 2020) through its regulation of genes including apolipoprotein genes (Sauvaget et al. 2002; Archer et al. 2005; Howell et al. 2014), the lipid transporter Cd36 (Martinez-Jimenez et al. 2010), and *Ppara* (Martinez-Jimenez et al. 2010; Chamouton and Latruffe 2012), many of which are linked to absorptive enterocyte differentiation (Chen et al. 2021). Previous studies identified reduced intestinal lipid uptake in *Hnf4a*^ΔIEC^ mice (Frochot et al. 2012), and reduced FAO and increased lipogenesis in *Hnf4ag^DKO^*, though these mice also have reduced enterocyte differentiation (Chen et al. 2019b; Chen et al. 2020). While loss of *Hnf4a* results in loss of *Ppara* expression and other FAO genes, loss or activation of *Ppara* does not result in differential expression of *Hnf4a* (**Fig. 4K**) (Bunger et al. 2007). We also find HNF4A promotion of enterocyte differentiation may expose a reservoir of lipid-liganded NR binding sites at the genes that regulate lipid catabolism and FAO that is suppressed by microbes, especially following a HFM (**Fig. 5B-D**). HNF4A therefore may potentiate a particular promotion of FAO genes by PPARA in enterocytes and IECs.

The identification of high HNF4A motif enrichment at GF+HFM relative to CV+HFM sites was coincident with the separation of H3K27ac sites high in GF+HFM with enterocyte regulatory regions and CV+HFM with ISC regulatory regions (**Figs. 4N,5B**). It is not clear what contributes to altered H3K27ac levels in our experiment, though TF candidates like HNF4A may recruit HDACs and histone acetyltransferases (Armour et al. 2017; Thakur et al. 2019). We are able to identify that enterocyte-specific enhancers are frequently HNF4A bound and preferentially activated in GF+HFM relative to CV+HFM. In contrast, ISC regulatory regions were not typically bound by HNF4A and were amplified in CV+HFM (**Fig. 5**). We see that isolated ISCs also show transcriptional differences between CV and GF conditions related to the crypt-villus axis (**Fig. 4J**). Although we also expected the entire block of enterocyte or ISC specific regulatory regions to behave similarly, we saw that the differential H3K27ac signals are amplified or muted depending on whether it is an HNF4A binding site in our IEC preparation.

Microbiota profoundly impact circadian rhythms, including through *Nfil3*, *Ppara*, and the histone deacetylase, *Hdac3* (Mukherji et al. 2013; Thaiss et al. 2016; Wang et al. 2017; Kuang et al. 2019). As our samples were collected at the same time of day, our experimental design provides limited insight into circadian processes. The lipid transporter CD36 has been proposed as a circadian-regulated, HDAC3-dependent promoter of lipid absorption (Kuang et al. 2019). Although a PPARA-activated gene, we generally find both H3K27ac and RNA-seq levels at *Cd36* are reduced by colonization and HFM in our experiment (**Supplemental Fig. 6L; Supplemental Table S2**) (El Aidy et al. 2013). Timing of food intake is also a contributor to circadian patterns, and addition studies are needed to further resolve how intestinal adaptation to acute and long-term dietary and microbial signals are integrated with circadian programs (Parkar et al. 2019).

## Methods

### Mouse IEC preparation for ChIP-seq RNA-seq and lipidomics

Mouse jejunal IEC preparations for down stream applications generated exactly as previously defined (Davison et al. 2017) and detailed with computational processing and additional experiments in **Supplemental material**.

## Data Availability

All raw and processed sequencing data generated in this study are available at the NCBI Gene Expression Omnibus (GEO; https://www.ncbi.nlm.nih.gov/geo/) under accession number GSE185835. Additional data under accession GSE90461.

## Acknowledgements

This work was supported by grants from the National Institutes of Health (R01-DK093399, P01-DK094779, R01-DK113123, R01-DK111857, R01 DK081426, P01-HL020948), as well as the Nuclear Receptor Signaling Atlas consortium (NURSA, U24-DK097748). Gnotobiotic mice were provided by the Center for Gastrointestinal Biology and Disease (P30-DK034987). We are grateful to Bonne M. Thompson and Joyce J. Repa for helpful advice on lipidomic analysis.

