## Supplemental Figures and Materials for "Transcriptional integration of distinct microbial and nutritional signals by the small intestinal epithelium"

**SUPPLEMENTAL FIGURES (below):**
**Supplemental Figure S1**: Lipidomic analysis of jejunal preparations.
**Supplemental Figure S2**: A diversity of transcriptional responses to colonization and HFM.
**Supplemental Figure S3**: Identifying putative transcriptional interaction genes.
**Supplemental Figure S4**: Identifying regulatory region response to colonization and HFM.
**Supplemental Figure S5**: Identifying putative interaction regulatory regions.
**Supplemental Figure S6**: HFM impacts different regulatory regions depending on a microbiota.
**Supplemental Figure S7**: Genes with multiple HNF4A binding sites are more often microbially suppressed.
**Supplemental figure S8**: IEC PPARA immunofluorescence in GF and CV.

**SUPPLEMENTAL TABLES (provided as separate spreadsheets):**
**Supplemental Table S1**: Lipidomics database
– Summary of output and comparisons from lipidomics of jejunal preparations in the GF, GF+HFM, CV and CV+HFM condition.

**Supplemental Table S2**: RNA-seq database
- Summary of DEseq2 output for RNA-seq comparisons across the GF, GF+HFM, CV, and CV+HFM conditions. RNA-seq LRT analysis and statistics is also included for each gene.

**Supplemental Table S3**: H3K27ac database
- Summary of DEseq2 output for significantly different H3K27ac ChIP-seq comparisons for enriched H3K27ac regions across the GF, GF+HFM, CV, and CV+HFM conditions.

**Supplemental Table S4**: H3K27ac LRT database
- Summary of DEseq2 output for H3K27ac LRT analysis across the GF, GF+HFM, CV, and CV+HFM conditions for enriched H3K27ac windows that pass defined significance thresholds.

**Supplemental Table S5**: HNF4A database
- Summary of DiffBind output for regions of merged HNF4A occupancy enrichment for comparisons across the GF, GF+HFM, CV, and CV+HFM conditions.

**Supplemental Table S6**: FAO genes
- Lists of FAO-related genes compiled from FAO-related GO terms.

**Supplemental Table S7**: Utilized published datasets
- Summary of utilized published datasets and processing.

**Supplemental Table S8**: PCR primers
- Summary of PCR primers used in this study.

### SUPPLEMENTAL METHODS

#### Mouse Husbandry

All mice used in this study were in the C57BL/6J strain originally sourced from Jackson Laboratories and maintained in the National Gnotobiotic Rodent Resource Center (NGRRC) at the University of North Carolina (UNC) at Chapel Hill. Male mice were reared under germ-free (GF) conditions, or reared GF and colonized (described below) with a conventional microbiota from C57BL/6J SPF mice for 14 d (conventionalized or CV). Mouse colonization was performed exactly as previously described (Davison et al. 2017). Production, colonization, maintenance, feeding, and sterility testing of GF mice were performed using the standard procedures of the NGRRC. Animals were housed on Alpha-dri bedding (Shepherd) and fed 3500 Autoclavable Breeder Chow (Prolab) or Picolab mouse diet 5058 (LabDiet) ad libitum. All experiments using mice were performed according to established protocols approved by the Institutional Animal Care and Use Committee at UNC at Chapel Hill.

#### BODIPY-egg yolk preparation and gavage

An egg yolk BODIPY labeled mixture was prepared as described with the following modifications (Carten et al. 2011): A 50% egg yolk mixture was generated using 2 ml of egg yolk (Grocery store bought) mixed with 2 ml PBS. The mixture was sonicated and strained as described to produce liposomes. 182 µl of BODIPY 558/568 $C_{12}$ (Invitrogen, D3835) was air dried and resuspended in 50µl of 100% ethanol and added to 2.3 ml of the 50% egg yolk liposomes mixture. 200 µl of mixture was used for gavage at 8:30 am EST.

#### Imaging and quantification of BODIPY labeled fatty acids in villi

Image analysis was blinded and performed using FIJI. Z-stack confocal images of fixed, flayed open, and whole-mounted duodenal intestinal segments were used for analysis. Z-stacks were taken across the axial plane of upward pointing villi. For each z-stack image, 3 planes were chosen for analysis, one near the top of the villus, one near the middle, and one towards the bottom villus. No two planes of each z-stack included the same cells. Villi were selected for analysis based on perfect or near-perfect vertical orientation to avoid analyzing cells from bent or angled villi. From each villus selected for analysis, 3 ROIs were defined which encapsulated about 5-15 epithelial cells but mostly excluded surrounding tissues or dark spaces. Sections of villi without clearly labeled nuclei were not selected as ROIs, as they could represent damaged or poorly imaged parts of the villus. For each villus quantified, one additional ROI within the same plane was taken of an empty space to be used as a baseline fluorescence value for normalization. Mean gray value of BODIPY-fatty acid (BODIPY-FA) conjugate was measured in each ROI. Mean gray value from nearest control ROI was subtracted from the mean gray value of each villus ROI. The resulting background subtracted mean gray values were averaged for each villus or mouse. 2 sample t-test was performed using JMP software to test for significant differences at both the per mouse and per villus level. Images with villus BODIPY-FA values closest to the average per-villus BODIPY-FA value for each group were selected as representative images.

#### PPARA Immunofluorescence and quantification

Mouse small intestinal (jejunum and ileum) tissue was dissected and immediately fixed overnight in zinc buffer formalin then embedded in paraffin. Five-µm thick paraffin sections were used for immunofluorescence experiments. Sodium citrate buffer (pH 6.0) was used for heat-induced antigen retrieval. Slides were blocked with 10% goat serum in PBS, then incubated with Rabbit Anti-PPARA (Invitrogen, PA1-822A) diluted 1:100 in antibody dilution buffer (PBS, 1% BSA, and 0.0025% Triton X-100) overnight at 4°C. Slides were washed with TBST (0.1% Tween-20), and incubated with Goat Anti-Rabbit Alexa Fluor 568 (Invitrogen, A-11011) diluted 1:200 in antibody dilution buffer. Slides were washed in TBST, counterstained with DAPI, and coverslips were mounted with ProLong Gold antifade reagent (Invitrogen, P10144). Slides were imaged on a Zeiss Axio Imager Z2 upright microscope with an apotome for optical sectioning. Nuclear PPARA fluorescence was quantified in the crypt and villus regions using ImageJ software. Two-way ANOVA (Prism) was used to analyze data (n = 4).

**Isolation of intestinal samples for RNA and ChIP**

Jejunal samples were collected, and processed, exactly as described (Davison et al. 2017). In our hands, these methods yield cell preparations that include most villus epithelial cells, a proportion of epithelial crypts, and a proportion of villus mesenchymal cells (data not shown). As the vast majority of cells in these preparations are epithelial, we operationally refer to them here as intestinal epithelial cells (IECs). CV and GF data for RNA-seq, DNase, H3K27ac, and HNF4A ChIP-seq was from Davison et al. 2017 (GSE90462) with the exception that two additional GF and two additional CV RNA-seq replicates were generated for this manuscript. All +HFM replicates were generated for this manuscript.

**RNA isolation, library preparation and sequencing**

Mouse jejunum IEC samples were subjected to RNA isolation, library preparation and sequencing as described (Davison et al. 2017). Briefly, prior to crosslinking, 1/50th of the isolated IECs were suspended in 1 ml TRIzol and stored at -80°C. Thawed IECs in

TRIzol were prepared according to the manufacturer directions: 200 μL of chloroform was added to the TRIzol and the sample was vortexed on high for 30 seconds at room temperature. The samples were incubated at room temperature for 2 minutes and centrifuged at 12,000 x g for 15 minutes at 4°C. The top aqueous layer was removed and added to equal volume of isopropanol. The nucleic acids were isolated using a column-based RNA-isolation kit (Ambion, 12183018A) with an on-column DNase I (RNase-free) treatment (New England Biolabs, M0303L) to remove DNA contamination. RNA was eluted in nuclease-free water, quantified using a Qubit 2.0 and stored at -80°C until submission to the Duke Sequencing and Genomic Technologies Core. RNA-seq libraries were prepared and sequenced by Duke Sequencing and Genomic Technologies Core on an Illumina HiSeq 2500 for 50 bp single end sequencing with 8 samples per lane in the flow cell.

**Chromatin immunoprecipitation, library preparation and sequencing**

Chromatin immunoprecipitation, ChIP libraries and sequencing was performed on mouse jejunum IECs as described (Davison et al. 2017). Briefly, frozen and sonicated chromatin from IECs was thawed on ice and diluted in 1 mL of ChIP dilution buffer (1% Triton X-100, 2 mM EDTA, 20 mM Tris-Cl (pH 8.1), and 150 mM NaCl) containing 1x Protease Inhibitor. This mixture was precleared with washed protein G Dynabeads (Thermo Fisher Scientific 10004D) for 3 hours at 4°C with gentle agitation. Beads were removed and chromatin was transferred to a clean microfuge tube and incubated with a ChIP-grade antibody [4 μg H3K27ac (Rabbit anti-H3K27ac, Abcam, ab4729), 8 μg HNF4A (Mouse anti-HNF4A, Abcam 41898)] overnight at 4°C with gentle agitation. Antibody-chromatin complexes were pulled down with washed protein G Dynabeads for 4 hours at 4°C with gentle agitation. The antibody-chromatin-bead complexes were washed 5x for 3 minutes with ice cold LiCl wash buffer (100 mM Tris-Cl (pH 7.5), 500 mM LiCl, 1% IGEPAL, 1% sodium deoxycholate) and 1x with ice cold TE buffer at 4°C on a nutator. Washed antibody-chromatin-bead complexes were resuspended in 100 µL of ChIP elution 12 buffer (1% SDS and 0.1 M sodium bicarbonate) and placed in a Eppendorf ThermoMixer C heated to 65°C and programed to vortex at 2000 RPM for 15 seconds, rest for 2 minutes for a total of 30 minutes. The beads were pelleted and the supernatant was moved to a new tube. This elution process was repeated once and corresponding elutions were combined for a total of 200 µL. To reverse crosslinked chromatin, 8 µL of 5 M NaCl was added to each 200 µL ChIP elution and was incubated at 65°C overnight. Immunoprecipitated chromatin was isolated using a QIAquick PCR quick preparation kit (Qiagen, 28104), quantified using a Qubit 2.0 fluorometer and stored at -80°C until library preparations and amplification. Libraries were always prepared within 3 days of the immunoprecipitation with the NEBNextUltra DNA Library Prep Kit for Illumina (New England Biolabs, E7370S). Prepared libraries were quantified using a Qubit 2.0 fluorometer and submitted to Hudson Alpha Genomic Services Laboratory for 50 bp single end sequencing on an Illumina HiSeq 2500 with 4 samples per lane in the flow cell. Germ-free or conventionalized chromatin for input normalization was generated using the same protocol as above except no antibody was used during the overnight antibody incubation; instead, chromatin was incubated at 4°C with gentle agitation. Bead incubation, reverse-crosslinking and library preparations for these samples were performed using the same protocol as the ChIPs.

**RNA-seq mapping and differential expression**

Adapter sequences and poor quality reads were removed from FASTQ files using trime_galore. Trimmed and high-quality sequences were aligned to the mouse genome (mm9) using STAR (version 2.7) using default parameters and length of genomic sequences around annotated junctions equal to 49. Counts per gene for differential expression analysis were generated using HTSeq (version 0.9.1) using default parameters. Differential gene expression analysis was performed using R (version 3.4.1) and DESeq2 (version 1.16.1). Counts from replicates for all 4 conditions were used to generate comparisons for interactions between nutritional status (-/+ high fat meal) and colonization status (-/+ microbes) using the Likelihood Ratio Test (test = "LRT") command from DESeq2. A cutoff of <.05 p-value and greater than 10 base mean counts was used to identify putative interaction genes. Individual comparisons (e.g. CV+HFM/GF+HFM) were extracted using the Wald test (test = "Wald") using an adj p-value of <.05 to define differential expression.

**ChIP-seq mapping, peak calls, and differential enrichment**

HNF4A, H3K27ac, and input GF+HFM and CV+HFM FASTQ sequences were mapped to mm9 genome using Bowtie2 version 2.3.2 and default parameters. MACS2 version 2.1.1.20160309 callpeak program was used to identify peaks from bowtie2-generated BAM files for H3K27ac and HNF4A ChIP samples using corresponding GF, GF+HFM, CV, CV+HFM, or input BAM files as a control, using the mouse mappable genome size (option: -g mm) and a bandwidth of 300 bp. Narrowpeaks output was utilized for HNF4A data. A single bed file of merged HNF4A ChIP peaks and a single bed file of merged H3K27ac ChIP peaks was generated by merging BED files from GF, GF+HFM, CV, and CV+HFM HNF4A replicates and GF, GF+HFM, CV, and CV+HFM replicates H3K27AC respectively using bedops v2.4.28 merge function. Blacklisted regions of the mm9 genome were removed from merged bed files using bedtools v2.26.0 subtract function. Merging peaks allowed for identifying changes for the same enriched regions across all 4 conditions and comparisons. To quantify counts for each merged H3K27ac peak for each replicate, sliding windows of 300 bp width and overlapping by 100 bp (200 bp steps) across H3K27ac merged peaks were generated using the R (v3.4.1) package

IRanges (v2.8.2) every 200 bp. featureCounts (v1.5.3) was used to generate counts for each H3K27ac window. Each 300 bp H3K27ac window was tested for LRT using counts from replicates for all 4 conditions to generate comparisons for interactions between nutritional status (-/+ high fat meal) and colonization status (-/+ microbes) using the Likelihood Ratio Test (test = “LRT”) command from DESeq2. A cutoff of less than 0.01 p-value and greater than 15 base mean counts was used to identify putative interaction regions. Individual comparisons (e.g. CV+HFM/GF+HFM) were extracted using DESeq2 using an adj p-value of <.05 to define differential enrichment. Significant windows of differential H3K27ac enrichment were then joined to merged H3K27ac peaks to allow for comparison across conditions. For HNF4A ChIP-seq, featureCounts (v1.5.3) was used to generate counts for each merged peak. DiffBind was used to generate differential peak calls for each comparison.

**DNase hypersensitivity site (DHS) peaks**

Merged jejunal IEC DHS peaks from CV and GF replicates were used from Davison et al. 2017 (GSE90462) (Davison et al. 2017).

**Transcription factor motif enrichment**

For each H3K27ac ChIP-seq directional-significance-group sequence from the closest jejunal DHS within 1kb of H3K27ac site was used for TF motif enrichment analysis. Input was composed of the associated DHS sequences for a single directional H3K27ac significance group. The background was composed of sequences from the reciprocal direction of the input group for the same comparison. TF motif enrichment was generated using the findMotifs command for Homer using vertebrate motifs with input and background as described for all 8 H3K27ac directional significance groups (Heinz et al. 2010).

**Definition of orthologs**

One to one mouse to human orthologs were extracted from Ensembl biomart (Ensembl genes 104) and were utilized to compare RNA-seq data from ileal Crohn's disease (Haberman et al. 2014) to mouse RNA-seq comparing expression levels in enterocytes versus ISCs (Kazakevych et al. 2017).

**Identification of neighboring genes for H3K27ac, HNF4A, and DHS**

To identify nearest neighboring genes (NCBIM37/mm9 Ensembl 91) for different regions of enrichment, ClosestBed was used with the first reported gene used for any ties. To quantify HNF4A sites per gene, each HNF4A site was assigned to a nearest gene and then the total HNF4A sites per gene was summed. For GREAT analysis, regulatory region coordinates were used as input using default parameters (McLean et al. 2010). GREAT may assign a region to multiple neighboring genes based on definitions that allow gene domains to overlap. These GREAT neighboring gene definitions were used only during reporting about GREAT analysis.

**Venn diagrams**

Venn diagrams were generated using an online Venn diagram maker (http://bioinformatics.psb.ugent.be/webtools/Venn/).

**FAO genes**

FAO genes were defined by extracting genes from FAO associated GO terms. The full list of genes and associations are listed in **Supplemental Table S6**.

**Kolmogorov–Smirnov**

To identify if a gene's differential expression level was significantly associated with a neighboring significantly differential H3K27ac site for each comparison a two-sided Kolmogorov-Smirnov test was performed in GraphPad Prism 9.

**GO term Metascape**

Differential RNA-seq gene lists were used as input for Metascape gene annotation and analysis (https://metascape.org) using default settings. For identification of coincident GO terms, the 8 RNA-seq directional significance gene lists were used as input using the multiple list meta-analysis option and output (Zhou et al. 2019).

**GREAT GO terms and site distribution**

Coordinates for the 8 H3K27ac ChIP-seq directional significance groups or other defined groups were used separately as input for GREAT (http://great.stanford.edu/public/html/index.php) using default settings (McLean et al. 2010). To generate coincident GO terms, exact matches for terms significant in both a +CV comparison and a +HFM comparison were displayed. Site distributions relative to transcriptional start sites were generated by using the 4 H3K27ac ChIP-seq comparison coordinates as input.

**Identification of motifs in accessible chromatin regions**

Genomic sequence from accessible chromatin regions from a merged set of peaks from IEC subtypes (Raab et al. 2020) was scored for HNF4A motif presence using the homer2 find command with the HNF4a(NR),DR1/HepG2-HNF4a-ChIP-Seq(GSE25021) position weight matrix. Motif identification in the *Ppara* regulatory region sequence was generated using the homer2 find command and vertebrate known motifs (Heinz et al. 2010).

**Additional dataset use**

When possible available datasets with fully genomewide results were utilized as provided in a publication's supplemental data or on GEO (https://www.ncbi.nlm.nih.gov/geo/). For incomplete microarray experiments, the GEO2R (https://www.ncbi.nlm.nih.gov/geo/geo2r/) analysis function was used with default settings and, unless specified a <.05 p-adj value was used as a cutoff for significance. For incomplete RNA-seq differential expression data, raw count files were used as input for DESeq2 or EdgeR with default settings. Data set use is summarized in **Supplemental Table S7**.

**Direct infusion MS/MS$^{ALL}$ lipidomic analysis**

Jejunal IECs were collected and isolated as described (Davison et al. 2017), and IEC pellets were snap frozen and stored at -80°C. IEC samples were transferred to a PTFE-lined screw-cap test tube containing 1 mL each methanol, dichloromethane, and water. Each mixture was vortexed and centrifuged at 3200 RPM for 5 min. The lower (organic) phase was transferred to a new tube using a Pasteur pipette and dried under $N_2$. The sample was reconstituted in 600 µL dichloromethane:methanol:isopropanol (2:1:1; v:v:v) containing 8 mM $NH_4F$ and 20 µL 3:50 diluted SPLASH® LipidoMix® internal standard. Using established methods (Vale et al. 2019), lipid extracts were infused into a SCIEX TripleTOF 6600+ mass spectrometer (Framingham, MA, USA) using a custom-configured LEAP PAL HTS-xt autosampler with dynamic load and wash (DLW) (Morrisville, NC, USA). Samples were infused into the mass spectrometer for 3 min at a flow rate of 10µL/min through the electrospray port of a DuoSpray ionization source. The MS/MS$^{ALL}$ data was obtained by acquiring production spectra at each unit mass between 200 and 1200 Da in positive mode. Electrospray voltage was set to 5500 V (and -4500 V in negative ionization mode), curtain gas (Cur) set to 20, Gas 1 and 2 set to 25 and 55 respectively, and temperature set to 300°C. Declustering potential (DP) and collision energy (CE) were set to 120V and 40eV for positive mode ionization. Collision energy spread (CES) function was not used as it resulted in incorrect isotope ratios which confounded the isotope correction algorithm in our software. Gas 1 and 2 as well as source gas were zero-grade air, and curtain gas and CAD gas was nitrogen. Processing the lipidomic data was done with in-house software (LipPY) developed at UT Southwestern Medical Center.

**Zebrafish husbandry**

Zebrafish lines were maintained using established protocols approved by the Office of Animal Welfare Assurance at Duke University. Conventionally raised zebrafish were reared and maintained as previously described (Westerfield 2000; University of Oregon Press). Production, colonization, maintenance and sterility testing of gnotobiotic zebrafish were performed as described (Pham et al. 2008).

**Cloning and Generation of *Tg(Mmu.Ppara:GFP)* transgenic zebrafish**

Construct-generation and generation of stable *Tg(Mmu.Ppara:GFP)* zebrafish on an EK zebrafish background was identical to those described previously using primers listed in **Supplemental Table S8**(Lickwar et al. 2017).

**Zebrafish imaging**

Whole mount 6 dpf zebrafish images were generated on a Leica M205 FA microscope with a Hamamatsu ORCA-Flash4.0 LT. 200 μm axial cross sections of the anterior intestine of 6 dpf crosslinked zebrafish were generated using a Leica VT1000S. Cross section images were generated using a Leica SP8 (DM6000CS) confocal microscope.

**Husbandry, gnotobiotic and HFM zebrafish treatment**

Generation, maintenance, and conventionalization of GF *Tg(Mmu.Ppara:GFP)* zebrafish larvae were conducted as described previously (Wen et al. 2021) with the exception that no exogenous food was administered until 6 dpf. At 6 dpf, half of the larvae of either GF or CV conditions were subjected to a high-fat meal as described in (Zeituni and Farber 2016), while the other half remained unfed. To feed a high-fat meal, larvae were transferred to sterile 6-well plates (20 larvae/well) and immersed in 5 ml solution of 5% chicken egg yolk liposomes in gnotobiotic zebrafish medium (GZM) for 6 h on a rocker at 28.5°C. After feeding, the fed larvae and their unfed counterparts were washed in GZM, euthanized, and collected in TRIzol for RNA isolation respectively (10-20 larvae per replicates; 5-6 replicates per condition).

**Preparation of zebrafish tissues and qRT-PCR**

mRNA was isolated from each replicate from pooled whole zebrafish larvae as previously described (Murdoch et al. 2019). RNA concentrations were measured using Thermo Scientific NanoDrop 1000 spectrophotometer and then diluted to match the lowest concentrated sample. The same total mRNA input (300 ng and 800 ng for the two independent experiments) was then used for cDNA synthesis using the iScript cDNA synthesis kit (Bio-Rad, 1708891). Quantitative real-time PCR was performed in triplicate for each replicate with 25 uL reactions using 2X Sybr Green SuperMix (PerfeCTa, Hi Rox, Quanta Biosciences, 95055) with the ABI Quantstudio 3 Real Time PCR system. Data were analyzed with the ΔΔCt method. Gene expression data was normalized using *ef1a* as the housekeeping gene. All statistical tests on qRT-PCR data were performed using Graphpad Prism v.9. Statistically significant effects of diet and microbial colonization on gene expression were determined by performing a 3-Factor RM ANOVA (gene x nutritional status x colonization) followed by a 2-Factor ANOVA for each gene. Post hoc unpaired 2-sided students' *t*-tests were applied to each gene exhibiting a significant nutritional status vs. colonization interaction.

### SUPPLEMENTAL FIGURES

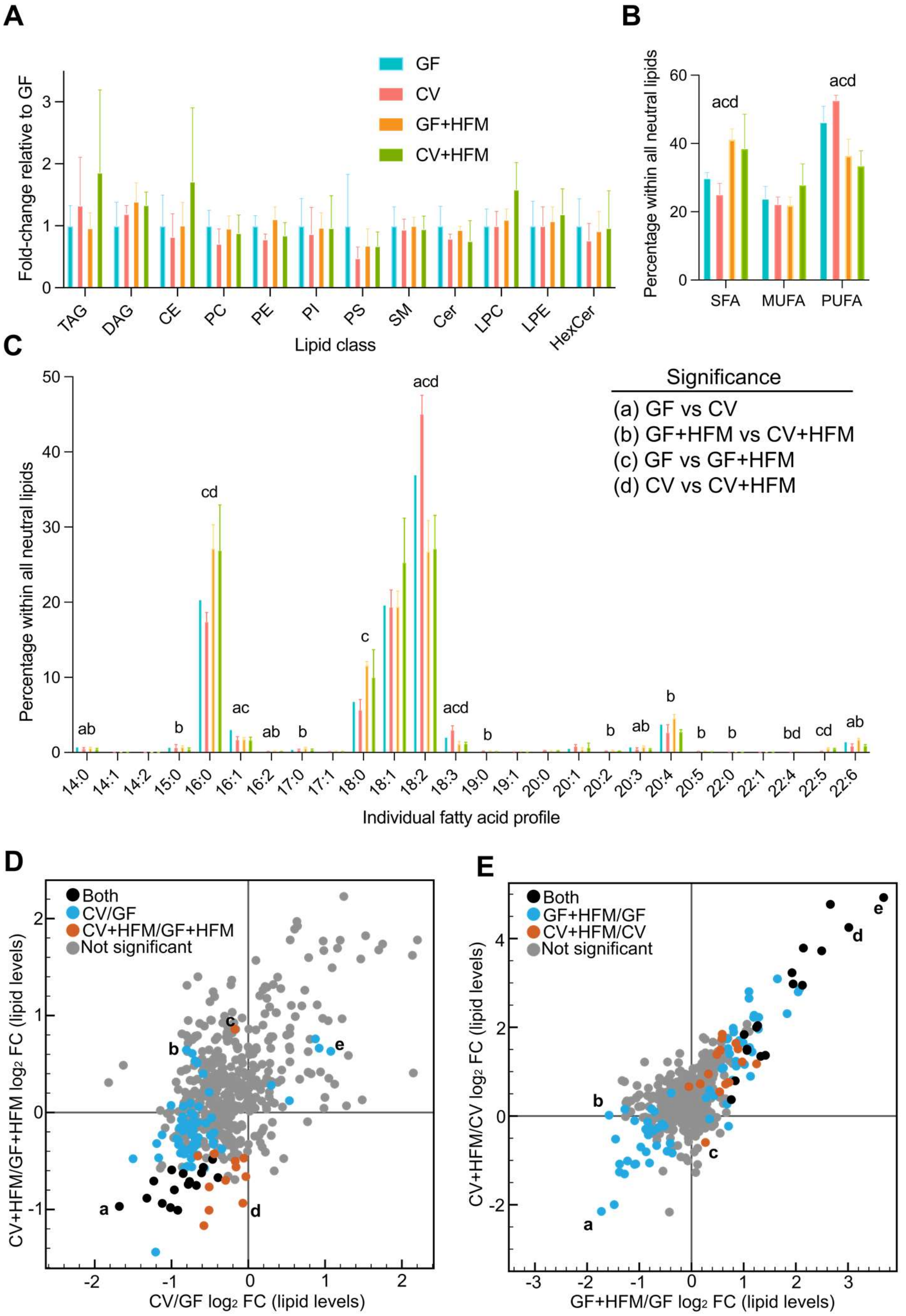


**a** PC|32:1|
**b** PC|35:5| or PC|36:4|(P-,)[H] or PC|36:5|(O-,)[H]
**c** PC|35:0| or PC|36:0|(O-,)[H] or PC|36:7| or PC|37:6|(P-,)[H] or PC|37:7|(O-,)[H]
**d** DAG|36:0|(NL-18:0,18:0)
**e** DAG|36:4|(NL-18:2,18:2)

**a** TAG|54:7|(NL-18:2)
**b** TAG|50:4|(NL-16:1)
**c** DAG|36:0|(NL-18:0)
**d** TAG|50:0|(NL-18:0)
**e** TAG|50:0|(NL-16:0)

**Supplemental Figure S1: Lipidomic analysis of jejunal preparations**. (**A**) Relative abundance of major lipid classes in each sample type including neutral lipids [triacylglyceride (TAG), diacylglyceride (DAG), cholesteryl ester (CE)], phospholipids [phosphatidylcholine (PC), phosphatidylethanolamine (PE), phosphatidylinositol (PI), phosphatidylserine (PS)], and polar lipids [sphingomyelin (SM), ceramide (Cer), lysophosphatidylcholine (LPC), lysophosphatidylethanolamine (LPE), hexosylceramides (HexCer)]. All measurements were normalized to internal standards and are shown as fold change relative to GF.

(**B**) Percentage of fatty acids detected within all neutral lipid classes that are saturated fatty acids (SFA), monounsaturated fatty acids (MUFA), and polyunsaturated fatty acids (PUFA).

(**C**) Percentage of fatty acids detected within all neutral lipid classes with the corresponding chain length and saturation. Data are shown as average and standard deviation of 4 mice per condition. Significant differences ($p<0.05$) by two-tailed Student's t-test comparing (a) GF vs CV, (b) GF+HFM vs CV+HFM, (c) GF vs GF+HFM, (d) CV vs CV+HFM are shown. See also Supplemental Table S1.

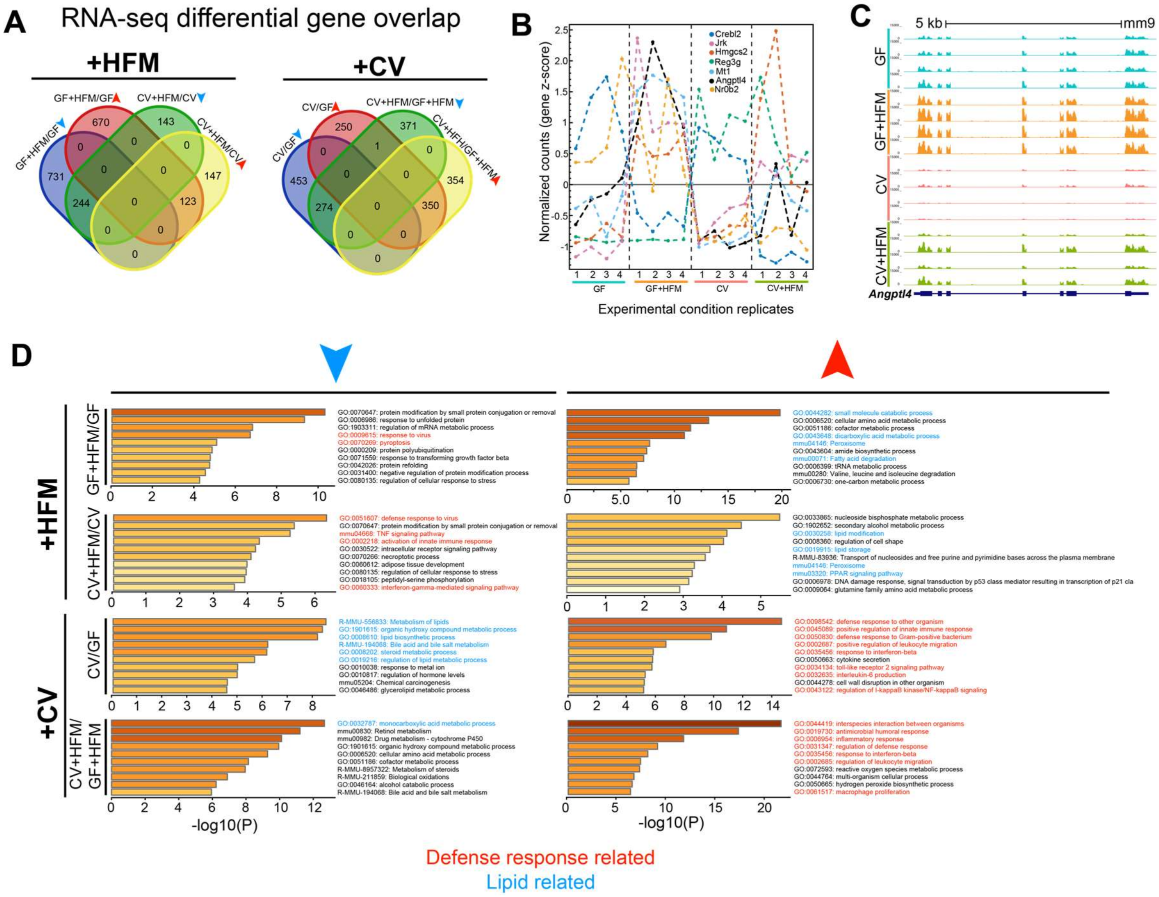


**Supplemental Figure S2: A diversity of transcriptional responses to colonization and HFM.**

**(A)** Venn diagrams for directional RNA-seq significance groups for +HFM and +CV comparisons.

**(B)** Example patterns of relative expression levels for genes significant in at least one comparison show diverse transcriptional regulation across the four conditions.

**(C)** UCSC screenshot for RNA-seq levels at the *Angptl4* locus.

**(D)** Metascape GO term enrichment for gene lists from each of the 8 directional RNA-seq significance groups. Defense response related terms are marked in red and lipid metabolism related terms are in blue.

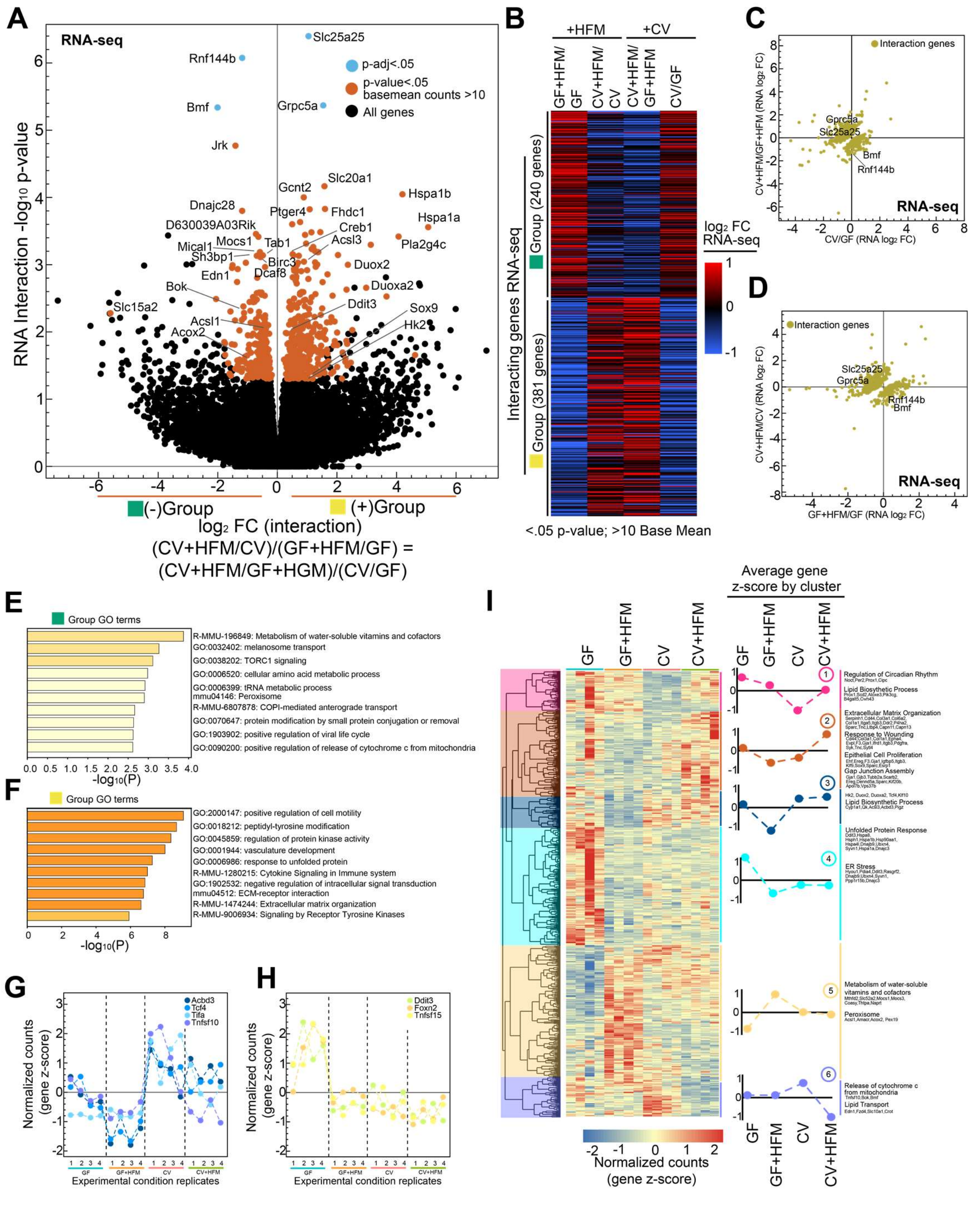


**Supplemental Figure S3: Identifying putative transcriptional interaction genes**

**(A)** Volcano plot showing interaction $\log_2$ fold change versus $-\log_{10}$ p-value for typical (p-adj<.05, >10 base mean counts; blue) and lenient (p-value<.05; red) cutoffs identifies genes with greatest potential for interaction. In effect, the interaction $\log_2$ fold change represents the $log_2$ ratio of (CV+HFM/CV)/(GF+HFM/GF) or (CV+HFM/GF+HFM)/(CV/GF) which are equivalent because these comparisons contain the same 4 conditions. Because negative (-, green) and positive (+, yellow) interactions are representative of the directionality of the fold change, but not necessarily the nature of the interactions, these groups are also colored to help illustrate that property.

**(B)** Heatmap of $log_2$ FC for each comparison for interaction genes broken into the green and yellow groups.

**(C)** Scatterplot of interaction genes comparing traditional $log_2$ fold change for CV/GF versus CV+HFM/GF+HFM RNA-seq.

**(D)** Same as C for GF+HFM/GF versus CV+HFM/CV.

**(E)** Metascape GO term output for green RNA interaction group genes.

(**F)** Metascape GO term output for yellow RNA interaction group genes.

**(G)** RNA-seq z-scored normalized counts for example interacting genes.

**(H)** RNA-seq z-scored normalized counts for example interacting genes.

**(I)** Clustered heatmap of gene z-scored normalized counts identify 6 clusters with average signal summarized in the middle and example genes and GO terms for each cluster to the right.

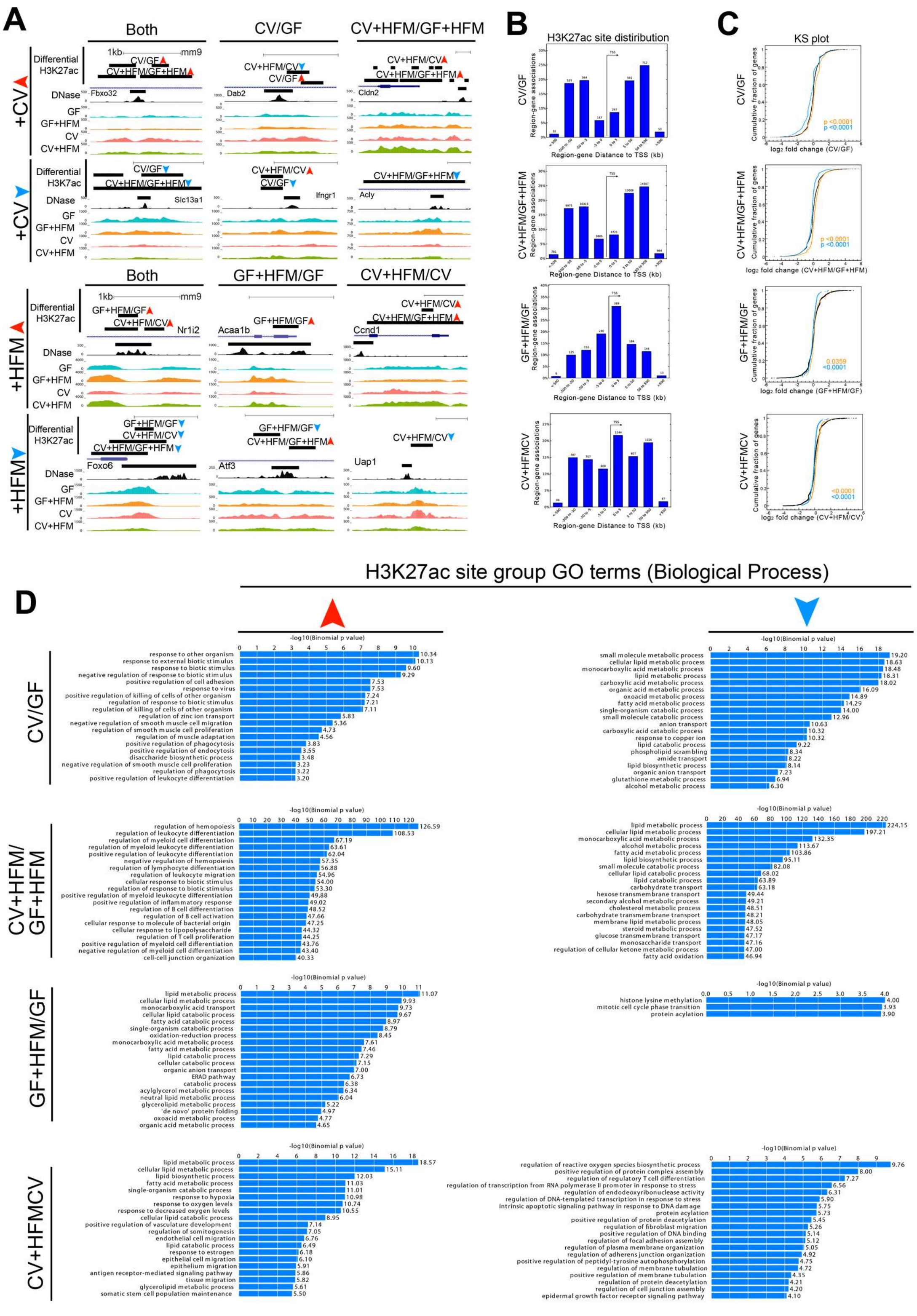


**Supplemental Figure S4: Identifying regulatory region response to colonization and HFM**

**(A)** Example loci showing significantly differential H3K27ac enrichment for various comparisons including across multiple comparisons.

**(B)** Distribution and number of sites relative to the nearest transcriptional start sites.

**(C)** Kolmogorov–Smirnov plots for each comparison shows transcript abundance of neighboring genes is positively associated with H3K27ac signal at differential sites.

**(D)** GREAT GO term enrichment using input of all sites for a particular group of differential H3K27ac sites.

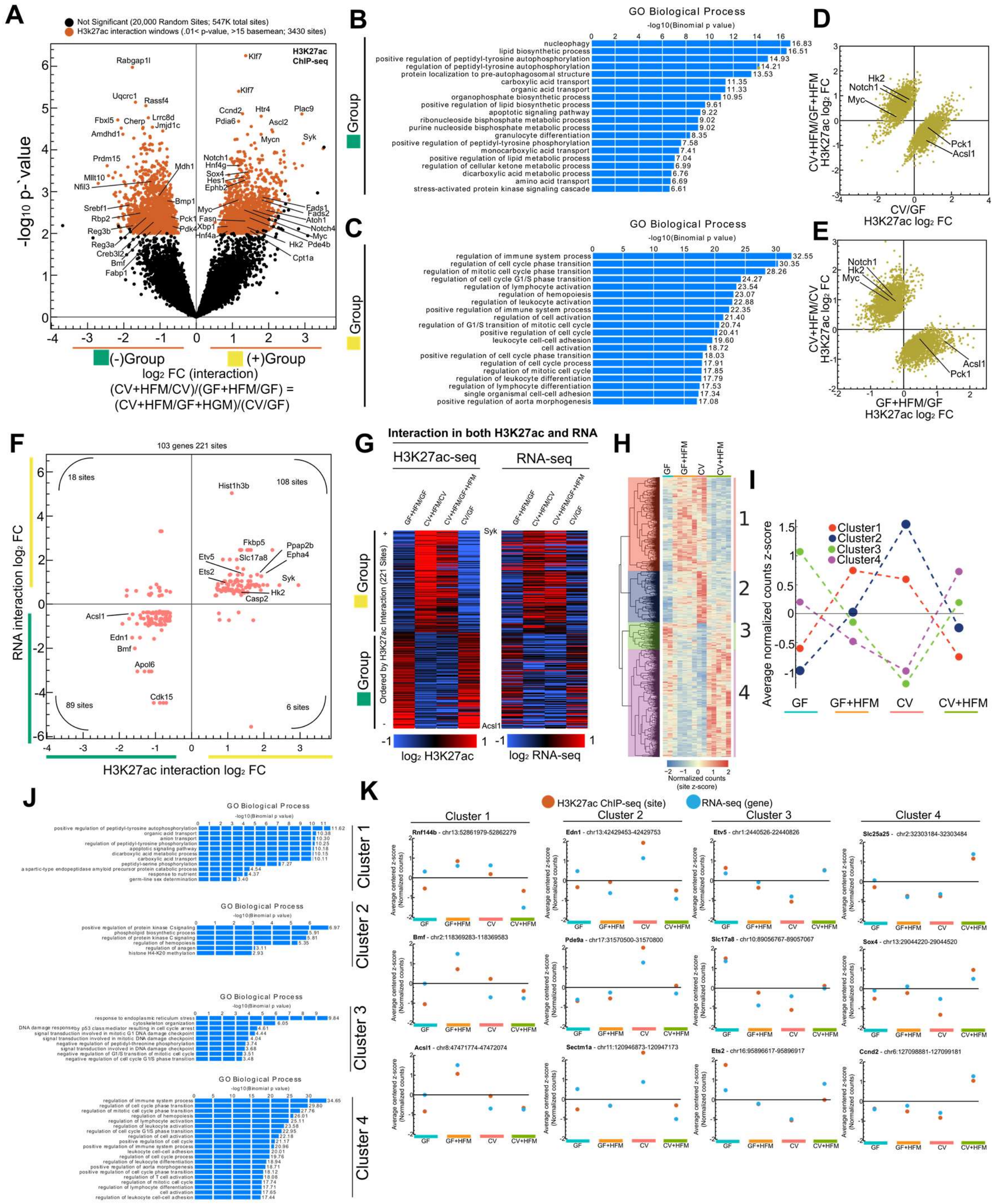


**Supplemental Figure S5: Identifying putative interaction regulatory regions**

**(A)** Volcano plot showing interaction $\log_2$ fold change versus $-\log_{10}$ p-value for lenient (p-value <.01, >15 base mean; red) cutoffs identifies H3K27acnregulatory windows with greatest potential for interaction. In effect the interaction $\log_2$ fold change represents the $\log_2$ ratio of (CV+HFM/CV)/(GF+HFM/GF) or (CV+HFM/GF+HFM)/(CV/GF), which are equivalent because these comparisons contain the same 4 conditions. Because negative (-, green) and positive (+, yellow) interactions are representative of the directionality of the fold change, but not necessarily the nature of the interactions, these groups are also colored to help illustrate that property. 20,000 out of 547,000+ enriched H3K27ac windows that did not pass the lenient interaction cutoff were chosen at random to represent non-interacting sites.

**(B)** GREAT GO term enrichment using input of the green interaction group of H3K27ac sites.

**(C)** GREAT GO term enrichment using input of the yellow interaction group of H3K27ac sites.

**(D)** Scatterplot of CV/GF versus CV+HFM/GF+HFM for H3K27ac interaction windows. Various genes allow for orientation to the volcano plot in (A)

**(E)** Same as (D) for GF+HFM/GF versus CV+HFM/CV.

**(F)** Scatterplot comparing H3K27ac $\log_2$ interaction windows fold change linked to the RNA $\log_2$ interaction fold change for genes that also show interaction reveals a positive correlation suggesting many of these regions are causal in contributing to the transcription patterns of these genes across +CV and +HFM conditions.

**(G)** Heatmap showing comparison of H3K27ac $\log_2$ fold change for all comparisons for green and yellow H3K27ac interaction groups show similar patterns of expression at linked gene neighbors.

**(H)** Clustered heatmap of z-scored normalized counts for H3K27ac interaction windows identify 4 clusters.

**(I)** Average normalized count z-score signal for the four clusters defined in (H).

**(J)** GREAT GO term enrichment for the H3K27ac clusters as defined in (H).

**(K)** Selected H3K27ac windows from each cluster showing consistent patterns of H3K27ac enrichment and relative RNA levels across the four conditions.

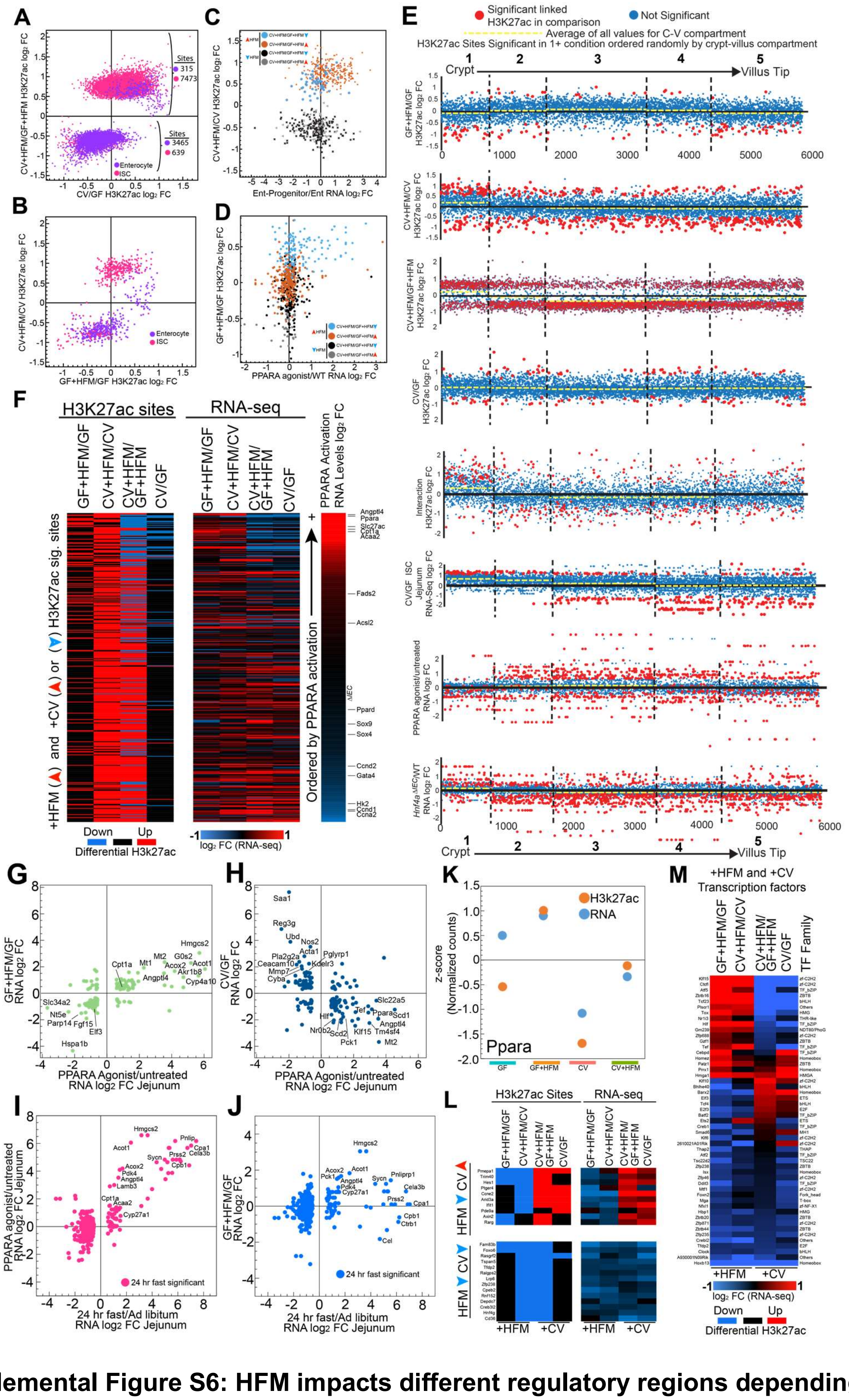

**Supplemental Figure S6: HFM impacts different regulatory regions depending on a microbiota**

**(A)** Scatterplot of significant H3K27ac sites for CV/GF and CV+HFM/GF+HFM $log_2$ FC colored by overlap with enterocyte or ISC regulatory regions.

**(B)** Same as (A) for GF+HFM/GF vs CV+HFM/CV

**(C)** Scatterplot of groups of H3K27ac sites that are significantly responsive in both a +CV and +HFM condition identifies that only sites that are +CV-up and +HFM-up are linked to genes that are expressed more highly in enterocyte progenitors as compared to enterocytes (Kim et al. 2014).

**(D)** Scatterplot of groups of H3K27ac sites that are significantly responsive in both a +CV and +HFM condition identifies that only sites that are +CV-down and +HFM-up are linked to genes that are activated by a PPARA agonist (Bunger et al. 2007).

**(E)** H3K27ac sites that are significantly different in at least one comparison grouped by their linked neighboring gene into one of five compartments based on preferential expression along the crypt-villus axis (Moor et al. 2018). Within the compartments H3K27ac sites are ordered by random. H3K27ac sites that are significant by the comparison on the Y-axis are red dots. All non-significant sites are blue. Yellow dashed lines are average for all sites within that compartment. Included are data sets from this study using H3K27ac $log_2$ FC for all the comparisons: GF+HFM/GF, CV+HFM/CV, CV+HFM/GF+HFM, CV/GF and the H3K27ac interaction analysis. Note the apparent strong difference in the behavior of CV+HFM/GF+HFM signal for sites along the 5 different zones, with zone 1 being elevated and zone 3 being the most reduced. This is consistent with differences in the crypt-villus axis described in Figure 5, but suggests preferential expression and transcriptional regulation along the villus may also contribute to differential signal. Also included are RNA data sets comparing CV/GF for Sox9+ sorted ISCs (Jejunum) (Peck et al. 2017), PPARA agonist/untreated (Jejunum) (Bunger et al. 2007), and *Hnf4a*$^{\Delta IEC}$/WT (Jejunum) (Verzi et al. 2013).

**(F)** Heatmap of +HFM-up H3K27ac that are also +CV-down or +CV-up across all comparisons ordered by linked genes $\log_2$ FC following PPARA activation/WT in descending order. Shown also are corresponding RNA-seq levels for linked genes and PPARA agonist/untreated $\log_2$ fold change levels (Bunger et al. 2007).

**(G)** Scatterplot of genes that are significantly differential following PPARA activation and in GF+HFM/GF show a positive correlation.

**(H)** Scatterplot of genes that are significantly differential following PPARA activation and in CV/GF show a negative correlation.

**(I)** Scatterplot of $\log_2$ fold change levels for genes significant (<.005 p-adj) in 24 hr-fast/ad-libitum versus PPARA activated genes (van den Bosch et al. 2007).

**(J)** Scatterplot of $\log_2$ fold change levels for genes significant (<.005 p-adj) in 24hr-fast/ad-libitum versus GF+HFM/GF shows many of the same PPARA targets are activated by fasting and a high fat meal (van den Bosch et al. 2007).

**(K)** z-scored normalized counts for the *Ppara* gene (RNA-seq) and *Ppara* H3K27ac site that is microbially suppressed and HFM induced and described in figure 3.

**(L)** Heatmap of example H3K27ac sites that are differentially enriched in +HFM-down as well as either +CV-down or +CV-up comparisons as well as have similar RNA-seq expression patterns. These are in contrast to the more highly characterized +HFM-Up and +CV-down or +CV-up sites.

**(M)** Heatmap of transcription factors that are significantly different in at least one +HFM and one +CV comparison by RNA-seq.

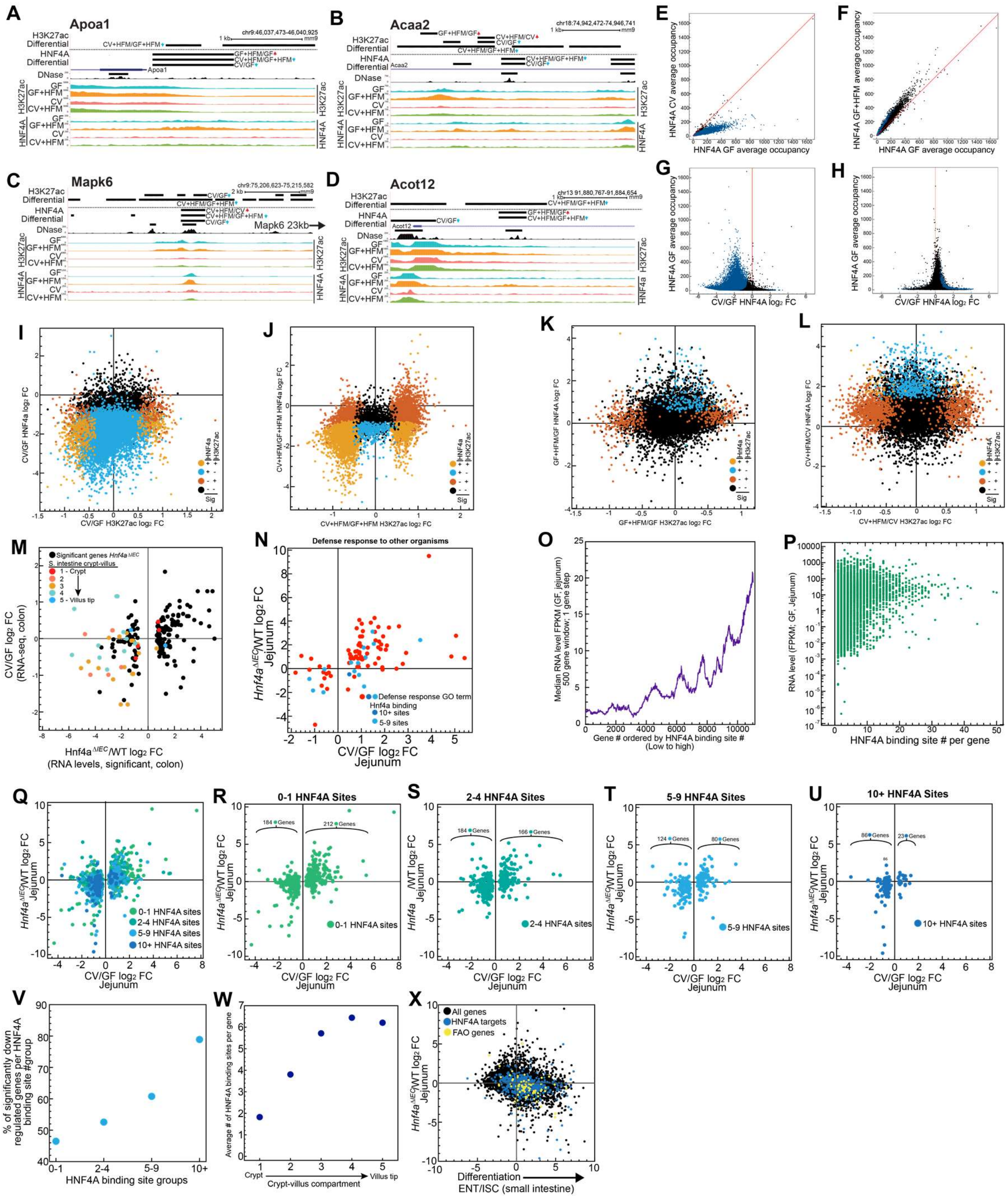


**Supplemental Figure S7: Genes with multiple HNF4A binding sites are more often microbially suppressed**

**(A)** UCSC screenshot of HNF4A binding sites that are significantly differential, corresponding to Figure 5B+C at the *Apoa1* **(A),** *Acaa2* **(B),** *Mapk6* **(C)**, and *Acot12* **(D)** loci.

**(E)** Scatterplot comparisons of HNF4A occupancy at all HNF4A binding sites (black) and significant sites (blue) GF versus CV HNF4A occupancy levels. Red line represents a slope of 1.

**(F)** Scatterplot comparisons of HNF4A occupancy at all HNF4A binding sites (black) and significant sites (blue) GF versus GF+HFM Hnf4a occupancy levels. Red line represents a slope of 1.

**(G)** Comparison of $\log_2$ HNF4A FC occupancy differences for HNF4A binding sites in the CV/GF comparison versus occupancy level in GF for all HNF4A binding sites (black) and sites significantly different in the CV/GF comparison (blue)

**(H)** Comparison of $\log_2$ FC occupancy differences for HNF4A binding sites in the GF+HFM/GF comparison versus occupancy level in GF for all HNF4A binding sites (black) and sites significantly different in the GF+HFM/GF comparison (blue).

**(I)** Scatterplot comparing CV/GF signal for overlapping H3K27ac and HNF4A enrichment sites colored by significance for each signal.

**(J)** Same as (I) for CV+HFM/GF+HFM

**(K)** Same as (I) for GF+HFM/GF)

**(L)** Same as (I) for CV+HFM/CV)

**(M)** Scatterplot comparing genes that are have significantly differential expression in *Hnf4a*$^{\Delta IEC}$/WT colon (Darsigny et al. 2009) versus CV/GF colon (Camp et al. 2014) colored by crypt-villus compartments from small intestine (Moor et al. 2018).

(**N**) Scatterplot of significant CV/GF $\log_2$ fold change RNA-seq levels versus *Hnf4a*$^{\Delta IEC}$/WT RNA levels for genes in the Defense Response GO term are not frequently bound by numerous Hnf4a binding sites.

**(O)** Genes ordered by the number of neighboring HNF4A binding sites show a positive correlation with gene expression level.

**(P)** Scatterplot showing the expression level of genes as the number of HNF4A binding sites gets high.

**(Q)** Scatterplot summary of significant CV/GF $log_2$ fold change RNA-seq levels versus *Hnf4a*$^{\Delta IEC}$/WT RNA levels colored by the number of neighboring HNF4A binding sites for each gene.

**(R)** Same as (Q) for 0-1 HNF4A binding sites per gene.

**(S)** Same as (Q) for 2-4 HNF4A binding sites per gene.

**(T)** Same as (Q) for 5-9 HNF4A binding sites per gene.

**(U)** Same as (Q) for 10+ HNF4A binding sites per gene.

**(V)** Percentage of significant CV/GF genes per HNF4A binding site group that are significantly down regulated for different binding site group bins.

**(W)** Average number of HNF4A binding sites per gene based on crypt-villus compartment groups (Moor et al. 2018).

**(X)** Scatterplot comparing small intestine RNA-seq $log_2$ fold change for Enterocyte/ISC shows a negative correlation with *Hnf4a*$^{\Delta IEC}$ suggesting genes with 10+ HNF4A binding sites (blue) contribute directly and indirectly to FAO genes (yellow) activation preferentially in enterocytes. Complement to Figure 4H.

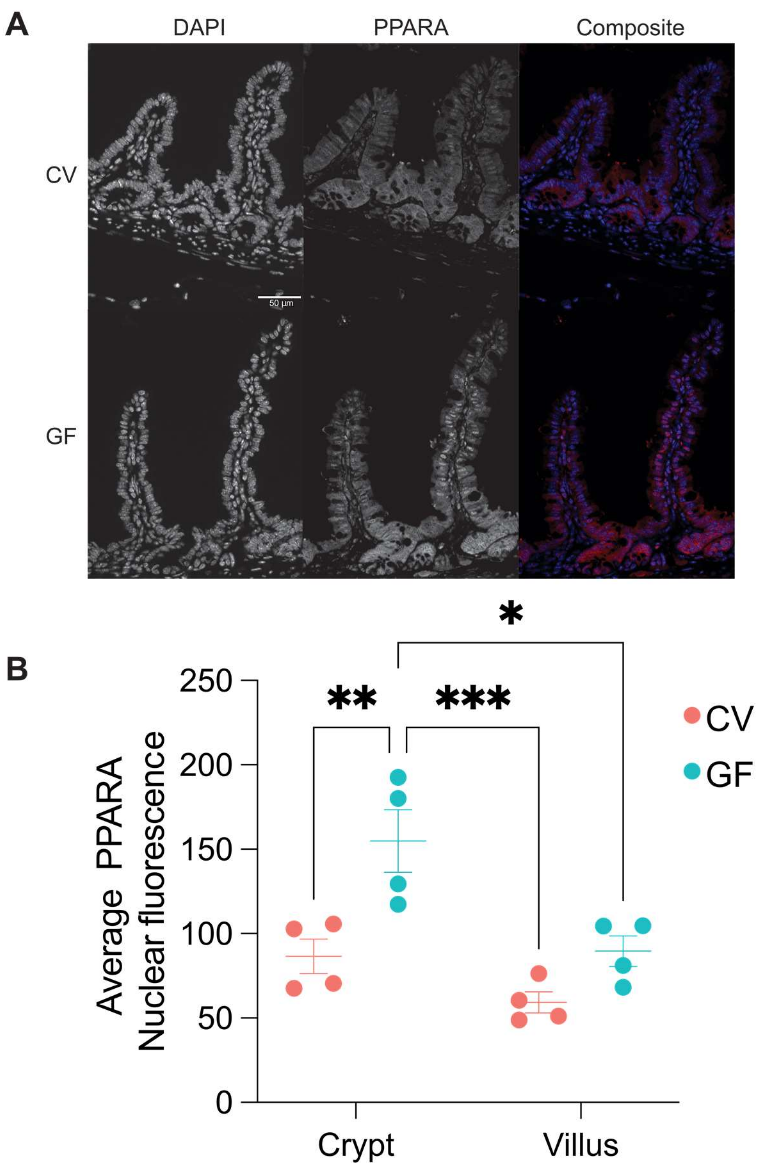


**Supplemental Figure S8: IEC PPARA immunofluorescence in GF and CV**

**(A)** PPARA immunofluorescence of small intestinal villi (red) in GF and CV conditions.

**(B)** Quantification of IEC PPARA nuclear fluorescence for the crypt and villus identifies higher nuclear fluorescence in GF. *p-value=<.05, **p-value=<.01, and ****p-value =<.0001 (n = 4 per condition). Anecdotally, in all conditions PPARA was higher in the cytoplasm in the crypt when compared to the villus.

Bunger M, van den Bosch HM, van der Meijde J, Kersten S, Hooiveld GJ, Muller M. 2007. Genome-wide analysis of PPARalpha activation in murine small intestine. *Physiol Genomics* **30**: 192-204.

Camp JG, Frank CL, Lickwar CR, Guturu H, Rube T, Wenger AM, Chen J, Bejerano G, Crawford GE, Rawls JF. 2014. Microbiota modulate transcription in the intestinal epithelium without remodeling the accessible chromatin landscape. *Genome Res* **24**: 1504-1516.

Carten JD, Bradford MK, Farber SA. 2011. Visualizing digestive organ morphology and function using differential fatty acid metabolism in live zebrafish. *Dev Biol* **360**: 276-285.

Darsigny M, Babeu JP, Dupuis AA, Furth EE, Seidman EG, Levy E, Verdu EF, Gendron FP, Boudreau F. 2009. Loss of hepatocyte-nuclear-factor-4alpha affects colonic ion transport and causes chronic inflammation resembling inflammatory bowel disease in mice. *PLoS One* **4**: e7609.

Davison JM, Lickwar CR, Song L, Breton G, Crawford GE, Rawls JF. 2017. Microbiota regulate intestinal epithelial gene expression by suppressing the transcription factor Hepatocyte nuclear factor 4 alpha. *Genome Res* **27**: 1195-1206.

Haberman Y, Tickle TL, Dexheimer PJ, Kim MO, Tang D, Karns R, Baldassano RN, Noe JD, Rosh J, Markowitz J et al. 2014. Pediatric Crohn disease patients exhibit specific ileal transcriptome and microbiome signature. *J Clin Invest* **124**: 3617-3633.

Heinz S, Benner C, Spann N, Bertolino E, Lin YC, Laslo P, Cheng JX, Murre C, Singh H, Glass CK. 2010. Simple combinations of lineage-determining transcription factors prime cis-regulatory elements required for macrophage and B cell identities. *Mol Cell* **38**: 576-589.

Kazakevych J, Sayols S, Messner B, Krienke C, Soshnikova N. 2017. Dynamic changes in chromatin states during specification and differentiation of adult intestinal stem cells. *Nucleic Acids Res* **45**: 5770-5784.

Kim TH, Li F, Ferreiro-Neira I, Ho LL, Luyten A, Nalapareddy K, Long H, Verzi M, Shivdasani RA. 2014. Broadly permissive intestinal chromatin underlies lateral inhibition and cell plasticity. *Nature* **506**: 511-515.

Lickwar CR, Camp JG, Weiser M, Cocchiaro JL, Kingsley DM, Furey TS, Sheikh SZ, Rawls JF. 2017. Genomic dissection of conserved transcriptional regulation in intestinal epithelial cells. *PLoS Biol* **15**: e2002054.

McLean CY, Bristor D, Hiller M, Clarke SL, Schaar BT, Lowe CB, Wenger AM, Bejerano G. 2010. GREAT improves functional interpretation of cis-regulatory regions. *Nat Biotechnol* **28**: 495-501.

Moor AE, Harnik Y, Ben-Moshe S, Massasa EE, Rozenberg M, Eilam R, Bahar Halpern K, Itzkovitz S. 2018. Spatial Reconstruction of Single Enterocytes Uncovers Broad Zonation along the Intestinal Villus Axis. *Cell* **175**: 1156-1167 e1115.

Murdoch CC, Espenschied ST, Matty MA, Mueller O, Tobin DM, Rawls JF. 2019. Intestinal Serum amyloid A suppresses systemic neutrophil activation and bactericidal activity in response to microbiota colonization. *PLoS Pathog* **15**: e1007381.

Peck BC, Mah AT, Pitman WA, Ding S, Lund PK, Sethupathy P. 2017. Functional Transcriptomics in Diverse Intestinal Epithelial Cell Types Reveals Robust MicroRNA Sensitivity in Intestinal Stem Cells to Microbial Status. *J Biol Chem* **292**: 2586-2600.

Pham LN, Kanther M, Semova I, Rawls JF. 2008. Methods for generating and colonizing gnotobiotic zebrafish. *Nat Protoc* **3**: 1862-1875.

Raab JR, Tulasi DY, Wager KE, Morowitz JM, Magness ST, Gracz AD. 2020. Quantitative classification of chromatin dynamics reveals regulators of intestinal stem cell differentiation. *Development* **147**.

Vale G, Martin SA, Mitsche MA, Thompson BM, Eckert KM, McDonald JG. 2019. Three-phase liquid extraction: a simple and fast method for lipidomic workflows. *J Lipid Res* **60**: 694-706.

van den Bosch HM, Bunger M, de Groot PJ, van der Meijde J, Hooiveld GJ, Muller M. 2007. Gene expression of transporters and phase I/II metabolic enzymes in murine small intestine during fasting. *BMC Genomics* **8**: 267.

Verzi MP, Shin H, San Roman AK, Liu XS, Shivdasani RA. 2013. Intestinal master transcription factor CDX2 controls chromatin access for partner transcription factor binding. *Mol Cell Biol* **33**: 281-292.

Wen J, Mercado GP, Volland A, Doden HL, Lickwar CR, Crooks T, Kakiyama G, Kelly C, Cocchiaro JL, Ridlon JM et al. 2021. Fxr signaling and microbial metabolism of bile salts in the zebrafish intestine. *Sci Adv* **7**.

Zeituni EM, Farber SA. 2016. Studying Lipid Metabolism and Transport During Zebrafish Development. *Methods Mol Biol* **1451**: 237-255.

Zhou Y, Zhou B, Pache L, Chang M, Khodabakhshi AH, Tanaseichuk O, Benner C, Chanda SK. 2019. Metascape provides a biologist-oriented resource for the analysis of systems-level datasets. *Nat Commun* **10**: 1523.
